# Transcriptional Dynamics of the Salicylic Acid Response and its Interplay with the Jasmonic Acid Pathway

**DOI:** 10.1101/742742

**Authors:** Richard Hickman, Marciel Pereira Mendes, Marcel C. Van Verk, Anja J.H. Van Dijken, Jacopo Di Sora, Katherine Denby, Corné M.J. Pieterse, Saskia C.M. Van Wees

## Abstract

The phytohormone salicylic acid (SA) is a central regulator of plant immunity. Antagonistic and synergistic actions between SA and other defense-associated hormones like jasmonic acid (JA) play key roles in determining the outcome of the plant immune response. To obtain a deeper understanding of SA-mediated transcriptional reprogramming and SA/JA crosstalk, we generated a high-resolution time series of gene expression from Arabidopsis leaves treated with SA alone and a combination of SA and methyl JA (MeJA), sampled at 14 time points over a 16-h period. We found that approximately one-third of the Arabidopsis genome was differentially expressed in response to SA, and temporal changes in gene expression could be partitioned into 45 distinct clusters of process-specific coregulated genes, linked to specific *cis*-regulatory elements and binding of transcription factors (TFs). Integration of our expression data with information on TF-DNA binding allowed us to generate a dynamic gene regulatory network model of the SA response, recovering known regulators and identifying novel ones. We found that 12% of SA-responsive genes and 69% of the MeJA-responsive genes exhibited antagonistic or synergistic expression levels in the combination treatment. Multi-condition co-clustering of the single- and combined-hormone expression profiles predicted underlying regulatory mechanisms in signal integration. Finally, we identified the TFs ANAC061 and ANAC090 as negative regulators of SA pathway genes and defense against biotrophic pathogens. Collectively, our data provide an unprecedented level of detail about transcriptional changes during the SA response and SA/JA crosstalk, serving as a valuable resource for systems-level network studies and functional plant defense studies.

## Introduction

In plants, like all complex organisms, the execution of specific gene expression programs is required to ensure proper development and appropriate responses to environmental perturbations and stresses. Phytohormones play important roles in regulation of gene expression programs. Direct and indirect interactions between genes and their molecular regulators, such as transcription factors (TFs), determine the complexity of the global large-scale reprogramming of gene activity (MacNeil and Walhout, 2011). Understanding the organization of these gene regulatory networks (GRNs) and how they ultimately drive specific biological outputs are key questions in plant biology, with the resulting knowledge important for driving improvement of agronomically important traits in crops (Ferrier et al., 2011; Krouk et al., 2013; Lavarenne et al., 2018).

To understand the dynamic organization and regulation of complex gene expression programs plant biologists are increasingly seeking to study systems in their entirety rather than focus on a small number of genes (Long et al., 2008). Such systems biology approaches are being driven by high-throughput omics technologies that can monitor different aspects of gene expression on a whole-genome scale. In particular, the accumulation of mature mRNA as determined by microarrays and high-throughput RNA sequencing (RNA-seq) analyses has emerged as the leading method for systems level gene expression profiling. Moreover, most biological processes are dynamic, and by collecting transcriptome data of high-resolution time series the regulatory networks and their sub-modules underlying gene expression programs can be more accurately deciphered. Several of these large-scale transcriptome studies have generated key biological and regulatory insights, including successful inference of GRN models for a wide-range of plant responses, including for example, development, hormone signalling, and abiotic and biotic stresses (Brady et al., 2007; Krouk et al., 2010; Breeze et al., 2011; Bechtold et al., 2016; Song et al., 2016; Hickman et al., 2017; Hillmer et al., 2017; Mine et al., 2018; Varala et al., 2018).

Phytohormones are key regulators of GRNs that are essential for a wide range of biological processes, including plant growth, development, reproduction, and survival. Changes in hormone concentration or sensitivity, as triggered among others by biotic stresses, induce gene expression programs that orchestrate a range of adaptive plant responses (Pieterse et al., 2012). Salicylic acid (SA) and jasmonic acid (JA) are major players in plant defense (Pieterse et al., 2012). While the JA pathway is generally induced by and effective against necrotrophic pathogens and herbivorous insects, the SA pathway is generally regulating resistance against (hemi)-biotrophic pathogens that feed and reproduce on living host tissue, such as *Pseudomonas syringae* and *Hyaloperonospora arabidopsidis* on *Arabidopsis thaliana* (hereafter Arabidopsis). SA has a well-established central role in the two major layers of plant immunity: pathogen-associated molecular pattern (PAMP)- triggered immunity (PTI), which is activated after pattern recognition receptors recognize conserved microbial patterns, and effector-triggered immunity (ETI), which is activated when the host detects perturbations in host cells caused by pathogen effector molecules (Vlot et al., 2009; Fu and Dong, 2013; Asai and Shirasu, 2015). SA is also involved in the broad-spectrum induced immune mechanism, known as systemic acquired resistance (SAR) (Klessig et al., 2018). Numerous studies, many of which utilized mutant screens, have revealed a diverse set of pathways, effector genes and host cellular processes as regulatory targets of SA (Vlot et al., 2009), including the production of antimicrobial pathogenesis-related (PR) proteins that promote immunity against diverse pathogens (Van Loon et al., 2006). In addition to its role in defense, SA also plays important roles in plant growth, development, thermotolerance, leaf senescence, responses to ultraviolet light and shade avoidance (Vlot et al., 2009; Nozue et al., 2018).

Over the past decade, studies in the reference plant Arabidopsis have dramatically enhanced our understanding of how SA is perceived and how this perception leads to changes in defense gene expression. Recent studies support a model in which the regulatory protein NONEXPRESSOR OF PR GENES1 (NPR1), and two close relatives, NPR3 and NPR4 each function as SA receptors (Fu et al., 2012; Wu et al., 2012; Ding et al., 2018; Innes, 2018). In the absence of a pathogen, when SA levels are low, NPR3- and NPR4-containing complexes are active and suppress the transcription of defense-related genes, while the activity of NPR1, which is a positive regulator of defense-related gene expression, is constrained at low levels of SA. Following pathogen detection, elevated SA levels and increased binding of SA by all three NPR proteins leads to a reversal in NPR activity. Binding of SA to NPR3 and NPR4 inactivates the co-transcriptional complexes, relieving their repressive activities (Ding et al., 2018), while NPR1 is activated by SA binding, leading to enhanced transcription of downstream target genes (Wu et al., 2012; Ding et al., 2018). Through interactions with TFs such as the TGA subclass of the basic leucine zipper (bZIP) family of TFs, NPR1 has been shown to be critical for the activation of a large fraction of SA-responsive genes, with a minority being regulated by an NPR1-independent mechanism (Wang et al., 2006).

Large-scale transcriptional reprogramming is a major feature of the SA response, and several studies have identified thousands of Arabidopsis transcripts that change in expression after treatment with either SA or benzothiadiazole S-methylester (BTH; a functional analog of SA) (Wang et al., 2006; Goda et al., 2008; Blanco et al., 2009), pointing to the existence of a complex GRN of TFs and target genes downstream of the core NPR-TGA transcriptional complexes. Several groups of TFs have been identified as regulators of the SA-mediated response. The role of members of the WRKY TF family in host immunity is firmly established, and several of them have been implicated in the regulation of SA-mediated transcriptional reprogramming by functioning as either positive or negative regulators of SA-responsive genes, including SA biosynthesis genes (Fu and Dong, 2013; Tsuda and Somssich, 2015). Selected members from other TF families, including NAC, TCP, and CAMTA have also been reported to regulate SA-responsive genes (Du et al., 2009; Zheng et al., 2012; Chandran et al., 2014; Wang et al., 2015; Zheng et al., 2015).

Different combinations of phytohormones coordinate the plant’s response to multiple simultaneous environmental inputs. Crosstalk between hormone signaling pathways generates an extra layer of complexity in the underlying GRNs, and allows for fine-tuning of adaptive responses in a cost-efficient manner (Spoel and Dong, 2008; Vos et al., 2013; Vos et al., 2015). In particular, the SA response is influenced by the antagonistic or synergistic action of other defense-associated hormones, like JA, abscisic acid (ABA), and auxin (Pieterse et al., 2012; Caarls et al., 2015). Antagonistic cross-communication between the SA and JA pathways, in both directions, is a paradigm for defense hormone pathway crosstalk, and has been demonstrated in several plant species (Pieterse et al., 2012). Attackers can exploit this antagonistic relationship to rewire the host immune response for their own benefit by the production of effector molecules that specifically activate either the SA or JA pathway to attenuate otherwise effective JA- or SA-mediated defenses, respectively (Brooks et al., 2005; El Oirdi et al., 2011; Gimenez-Ibanez et al., 2014). Recent studies have begun to provide insights into the mechanisms underlying crosstalk between the SA and JA pathways. The ERF TF ORA59, which positively regulates JA-signalling, had reduced transcript and protein levels upon activation of the SA pathway (Van der Does et al., 2013; Zander et al., 2014). The SA response was shown to be antagonized by three closely related JA-induced NAC TFs, which affected SA biosynthesis and metabolism (Zheng et al., 2012).

Despite the major role of the SA pathway in mediating plant immunity, relatively little is known about the organization of the GRNs that operate downstream of SA perception and the NPR signaling module, including which members from the diverse TF families orchestrate the elaborate gene expression programs that target distinct biological processes and lead to SA-mediated immunity. Moreover, the dynamic interplay between the SA and JA pathways remain poorly understood, with little knowledge regarding the genes and associated biological processes targeted, the kinetics of these interactions, and the underlying regulatory mechanisms,. Previously, we used high temporal resolution transcriptome profiling of MeJA-treated Arabidopsis leaves, which generated a series of novel insights into the chronology and regulation of the JA response and identified novel players in JA-mediated defenses to caterpillars and necrotrophic pathogens (Hickman et al., 2017). In the present study, we profiled the transcriptional dynamics in Arabidopsis leaves in response to SA and a combination of SA + MeJA to (1) obtain a detailed understanding of the architecture and dynamics of the GRN that underlie the SA response, (2) examine the dynamic nature and extent of crosstalk between SA- and JA-response pathways at the transcriptional level, and (3) predict and validate novel regulators of the SA-controlled GRN and SA-mediated immunity.

## Results

### A Time Course of SA-Elicited Transcriptional Reprogramming

We performed high-resolution transcriptional profiling of the SA response in Arabidopsis by using RNA-seq to measure mRNA levels in leaves harvested at 14 consecutive time points (15 and 30 min, and 1, 1.5, 2, 3, 4, 5, 6, 7, 8, 10, 12 and 16 h) following treatment with SA or a mock control. True leaf 6 from four independent 5-week-old Col-0 plants that were rosette-dipped in the SA or mock solution, were sampled in quadruplicate, providing four biological replicates per time point, per treatment. This leaf tissue was treated and harvested simultaneously with the material used in our previously published time series study of the JA response (Hickman et al., 2017). The RNA-seq libraries preparation and sequencing was performed in parallel with that of the JA study, and therefore the mock-treated control samples are shared between the two studies. To identify genes that responded to SA, we fitted a generalized linear model (GLM) to our transcription data, which led to the identification of 9524 differentially expressed genes (DEGs) between SA and mock treatments over the time course (Supplemental Data Set S1; see Methods). Many of these genes were not previously described as SA responsive in earlier array-based studies, where one to three time points were assayed in seedlings treated with SA (Goda et al., 2008; Blanco et al., 2009) or whole plants treated with benzothiadiazole S-methylester (BTH; a functional analog of SA) (Wang et al., 2006). Moreover, in our experiment the SA treatment clearly had a greater effect on the overall gene expression than MeJA treatment that changed the expression of 3611 genes (Hickman et al., 2017).

Inspection of the expression profiles revealed a variety of dynamic expression patterns, including genes that showed transient responses, sustained induction/repression, and some with complex behavior. Next, we analysed the temporal dynamics of the SA-response by estimating the time at which each gene first became differentially expressed using a previously outlined strategy (Hickman et al., 2017). Genes were subsequently divided into two sets of up- and downregulated DEGs (based on agglomerative hierarchical clustering, described in greater detail below). Plotting of the time point of first differential expression for all up- and downregulated DEGs indicated key time points of transcriptional change, which pointed to distinct transcriptional waves (Figure 1). Three waves of upregulation and two waves of downregulation were identified. For both gene sets the first wave of expression (0–2 h after SA treatment), representing the immediate transcriptional response to SA, contains a far greater number of DEGs than the subsequent waves, representing the intermediate/late SA response.

**Figure 1.**
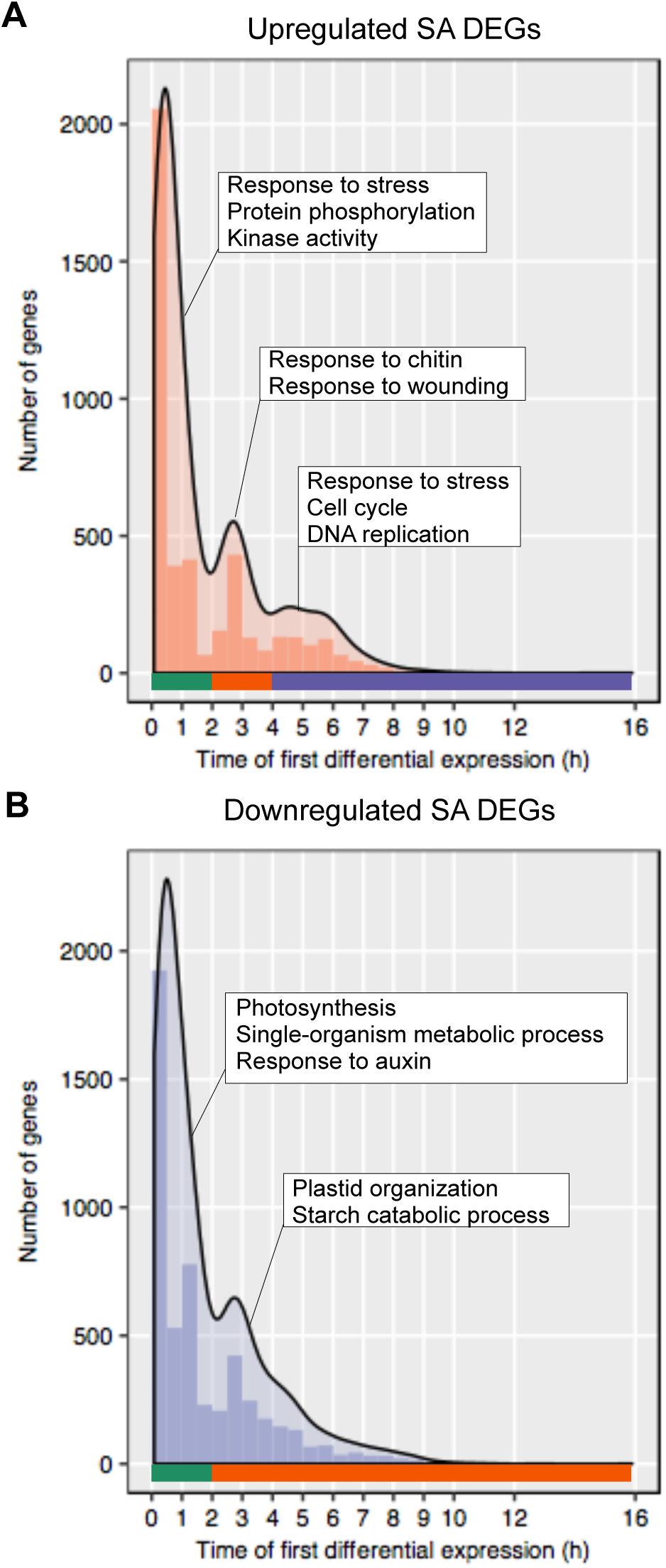
Major transcriptional waves of the SA response. Shown is the number of genes estimated to be first differentially expressed in 30 min time windows for genes upregulated **(A)** and downregulated **(B)** in *Arabidopsis* leaves following exogenous application of SA. Colored bars indicate waves of transcription, with green, orange and purple indicating the first, second and third waves respectively. Transcriptional waves are annotated with the most significant functional categories in each wave together with additional selected significant terms.

To investigate the biological relevance of these transcriptional waves, the genes assigned to each wave were tested for overrepresented functional categories using Gene Ontology (GO) term enrichment analysis. Interestingly, we found that each wave was enriched for distinct annotations, indicating a chronology in the regulation of different biological processes mediated by SA signalling. The first and largest wave of gene upregulation (Figure 1A) was linked to immune signalling, such as ‘protein phosphorylation’, ‘signal transduction’, ‘vesicle mediated transport’ and ‘programmed cell death’. Among the genes upregulated in the second wave (2–4 h) were some enriched for JA-related functional terms such as ‘response to chitin’ and ‘response to wounding’. The third, and final, wave of upregulation (4 h onwards) was associated with cell cycle and DNA repair, with the latter process previously reported as a target of SA signalling (Yan et al., 2013). Both waves of downregulated gene expression by SA (Figure 1B) were overrepresented for terms related to photosynthesis and include genes related to photosystem I and II, and chlorophyll biosynthesis. Distinct term enrichment was also apparent; for example, the terms ‘response to auxin’ and ‘starch metabolism’ were selectively enriched in the first and second waves, respectively. Concluding, this analysis showed that distinct pathways become active at different times during the SA response, indicating that these time series data possesses necessary information for delineating the dynamic use of transcriptional networks induced by SA.

### Coherent Functional Modules of Coexpressed Genes

To identify tightly regulated groups of coexpressed genes during the SA response, we used the time-series clustering algorithm SplineCluster to group the 9524 DEGs into 45 clusters of coexpressed genes that share similar expression profiles (Figure 2A; Supplemental Data Set S2). To investigate the biological significance of the distinct dynamic expression patterns, the genes in each cluster were tested for overrepresented functional categories using GO term enrichment analysis (Supplemental Data Set S2). Many of the clusters showed enrichment for coherent functional annotations and showed a wide-range of distinct biological processes to be regulated by SA (Figure 2B).

**Figure 2.**
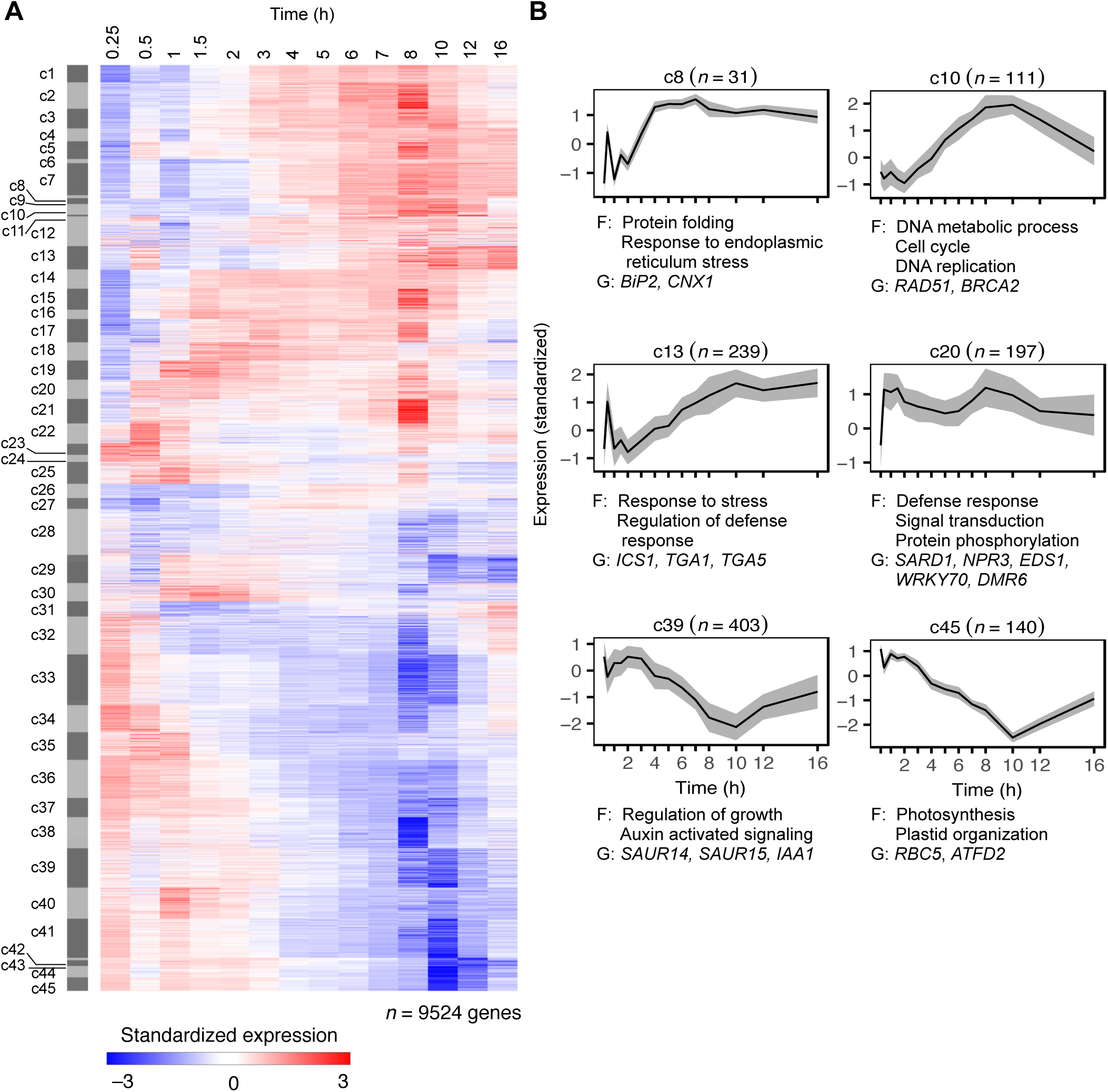
Temporal analysis of SA-responsive genes reveals coherent functional modules of coexpressed genes. **(A)** The set of 9524 genes showing differential expression in response to SA were partitioned into 45 (c1–c45) distinct coexpressed gene clusters using SplineCluster. Each row of the heatmap represents an individual gene and indicates level of expression, with red and blue indicating increased and decreased expression (standardized on a per gene basis across mock and SA treatments), respectively. **(B)** Representative coexpression modules with distinct functional enrichment. Shown is the mean expression profile for a module (black line), with grey areas indicating median absolute deviation. Cluster number and size (*n*) is indicated above each plot. Selected overrepresented functional categories (F) and representative genes (G) are denoted.

As expected, several upregulated coexpression clusters were enriched in immune-related annotations (e.g., c13, c19, c20), and correlate with many previously described primary SA-responsive genes, including genes involved in SA metabolism (*ICS1, PBS3, EDS5, DRM6, CBP60g, and SARD1*), and known regulators of the SA pathway (*TGA1*, *TGA5*, *NPR4*, *NIMIN1, NIMIN2, WRKY18*, *WRKY51*, *WRKY54* and *WRKY70*). Some coexpression modules were enriched for other defined SA-regulated biological pathways and processes. For example, c8 was associated with the protein folding response (Wang et al., 2005), and c10 was highly enriched for genes involved in DNA replication and cell cycle regulation (Wang et al., 2010; Wang et al., 2014).

Downregulated clusters showed how SA may regulate the switch from normal growth to defense with broad overrepresentation for functional terms associated with growth and development, and primary metabolism. Strikingly, there was strong enrichment of genes associated with photosynthesis among downregulated genes. Clusters c41–c45, all of which showed a marked decrease in transcript levels following SA treatment, were overrepresented for photosynthesis and plastid-related genes. Distinct enrichment of genes associated with development was also evident; for example, c35 and c39 are specifically overrepresented for genes associated with cell wall organization/biogenesis and auxin signaling, respectively. SA-mediated suppression of the auxin pathway has been shown to be an important component of plant defense (Wang et al., 2007).

Thus, our information-rich time series expression profiles revealed in unprecedented detail the broad range of biological processes differentially regulated by SA, and refines the breadth of the SA-responsive gene expression program. The identification of tightly-regulated groups of genes and the specific and coherent functional enrichment found in both up- and downregulated gene clusters suggests an elaborate underlying GRN, which controls these dynamic and varied changes in gene expression and biological processes to ultimately favor defense over normal growth and development.

### Transcriptional Control of the SA Response

To investigate mechanisms of gene regulation underlying SA-mediated transcriptional reprogramming, we analysed the coregulation of transcriptional response of Arabidopsis to SA at two levels of granularity. First, we created a set of coarse-grain SA-response modules, which we defined as the unions of upregulated (c1–c25) and downregulated (c26–c45) gene clusters and analyzed these gene sets for enriched TF-families and *cis-*regulatory elements derived from protein-binding microarray studies (Franco-Zorrilla et al., 2014; Weirauch et al., 2014), in order to capture broad mechanisms of coregulation.

Within the SA-upregulated set, genes encoding members of the WRKY and NAC TF families were significantly overrepresented (Figure 3A). In line with the enhanced activity of genes encoding these TFs, motifs corresponding to DNA binding sites of WRKY (W-box) and NAC TFs were also overrepresented in the group of upregulated genes, suggesting that these TF families dominate the onset of SA-induced gene expression (Figure 3B). While the genes encoding members of the bHLH and bZIP TF families are not enriched among SA-upregulated genes, the enrichment for their DNA recognition patterns, which consist of the core G-box motif, suggests a significant role for members of these families in the regulation of the SA response (Figure 3B). Genes in the downregulated coarse-grained module also displayed coherent enrichment for TF-family genes and their cognate DNA binding motifs. This set of SA-repressed genes was overrepresented for members of the CO-like, HD-ZIP, and bHLH TF-families (Figure 3A), with binding motifs for the HD-ZIP and bHLH families also enriched (Figure 3B). TFs belonging to these families have been shown to regulate processes associated with development, photosynthesis, and JA-dependent responses (Ariel et al., 2007; Leivar and Quail, 2011; Gangappa and Botto, 2014).

**Figure 3.**
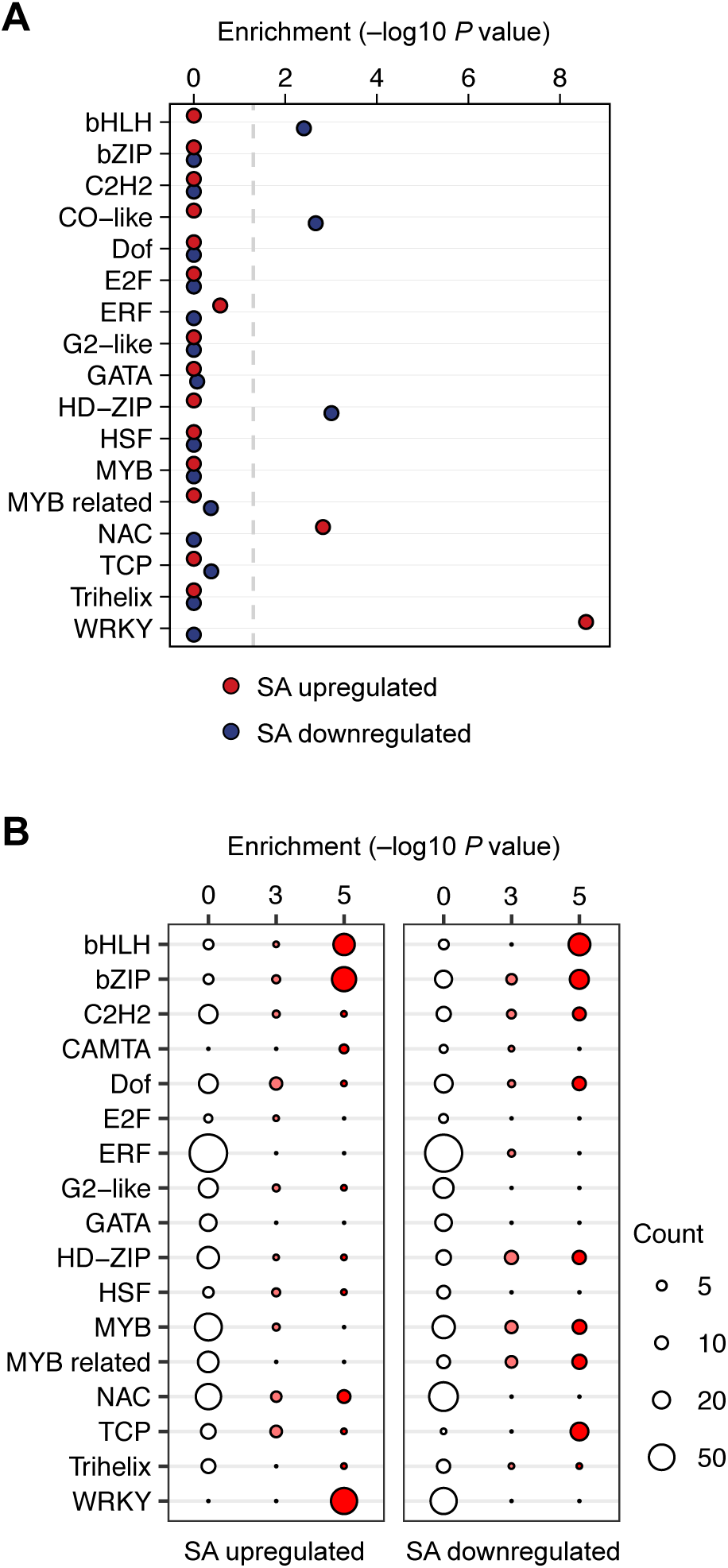
Enriched TFs and DNA-binding motifs in genes upregulated and downregulated in response to SA. **(A)** Significantly overrepresented genes of TF families within the unified upregulated (c1–25; red) or downregulated (c25–45; blue) gene clusters in response to SA treatment. Corrected *P* value < 0.01 (indicated by dashed grey line) (hypergeometric test). **(B)** Significantly overrepresented DNA-binding motifs in the promoters of genes upregulated or downregulated in response to SA treatment. Motifs are grouped according to cognate TF family. The size and colour of each circle represent the per-family motif count and enrichment *P* value range (hypergeometric test), respectively. Significantly enriched motif frequencies are indicated by red colouring.

Next, we investigated more specific regulatory mechanisms that may control the smaller subsets of process-specific SA-responsive genes, by analyzing motif enrichment within each of the 45 fine-grained coexpression modules identified by the SplineCluster analysis (Figure 4). This revealed promoter motifs that were selectively enriched in specific clusters, and offered a more precise link between motifs and module-specific expression patterns. DNA-binding motifs associated with WRKY TFs were highly enriched in many upregulated coexpression clusters, further demonstrating the importance of this TF family and its cognate binding sites in the regulation of the SA response. Binding sites for NAC TFs were also significantly and selectively overrepresented in a number of upregulated clusters. For both the WRKY and NAC DNA-binding motifs, the broad pattern of enrichment is consistent with the large-scale induction of genes that encode TFs from these families following SA treatment (Figure 3A). The motif analysis also highlighted several upregulated clusters that were not overrepresented for WRKY-binding motifs, suggesting additional TFs and *cis*-elements primarily regulate genes in these groups. For example, c9 and c10 are enriched for E2F TF DNA-binding motifs, while c12 and c22 are enriched for binding sites for HSF and CAMTA TF families, respectively. Consistent with the functional term analysis results, TF families implicated with regulation of growth and development are enriched in several downregulated clusters. For example, TCP and HD-ZIP TF DNA-binding motifs are selectively enriched in downregulated clusters, and members of both of these families are linked with regulation of growth and development (Martín-Trillo and Cubas, 2010). Binding sites for bHLH and bZIP TF families (both of which can bind DNA sites that contain the core G-box motif) were enriched in both selected upregulated and downregulated clusters, suggesting that G-box motifs may play an important role in directing both the up- and downregulation of target genes as part of the SA response.

**Figure 4.**
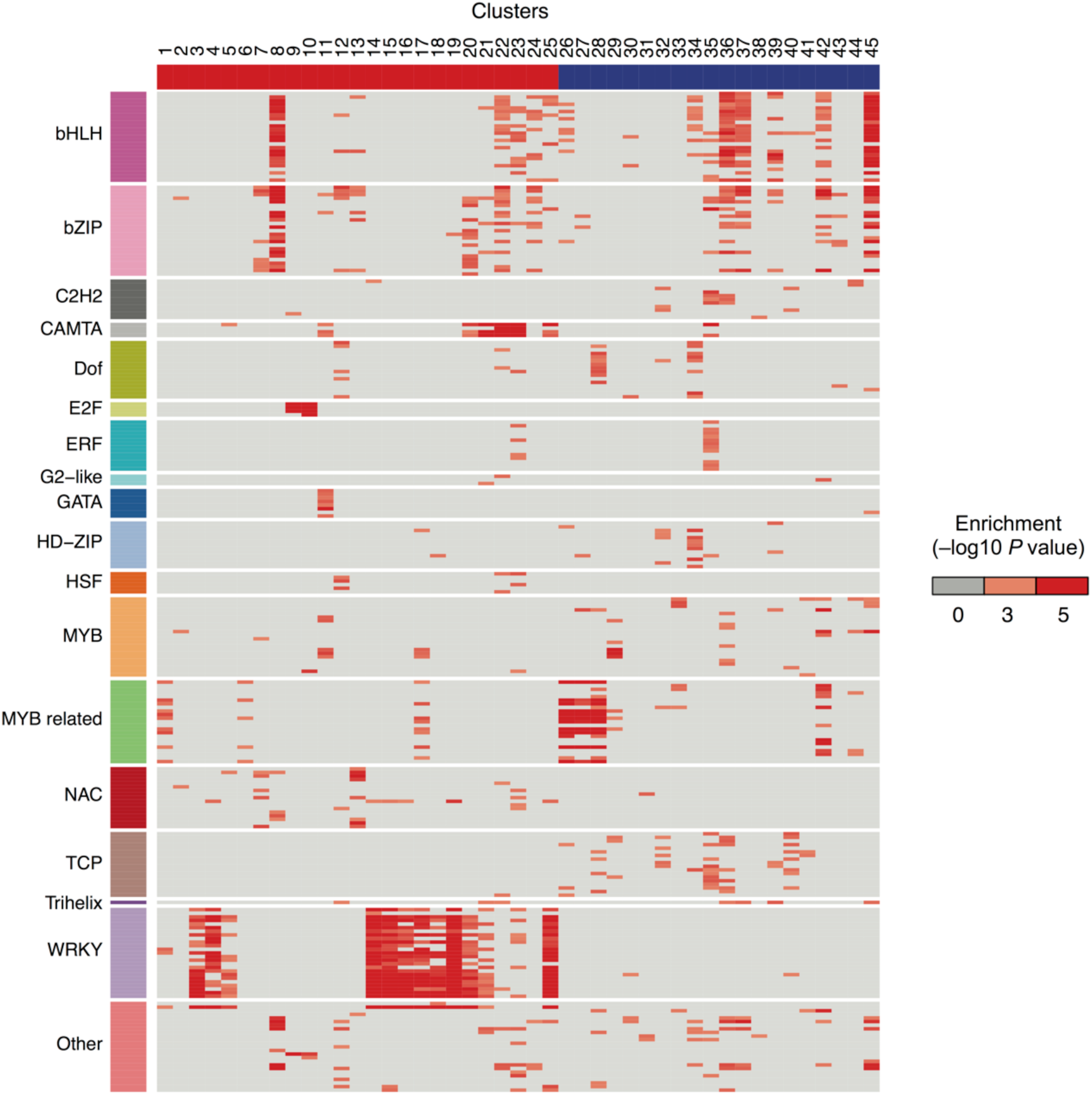
Enriched *cis*-regulatory motifs in SA-responsive gene coexpression clusters. Known TF DNA-binding motifs are differentially enriched in the promoters of genes clustered on the basis of their expression following application of SA. Rows indicate motifs and are colored by corresponding TF family. The top-ranked 25 motifs (by enrichment *P* value) are shown per family. Columns indicate co-expression clusters, with red and blue column colors differentiating between up- and downregulated clusters respectively. Red boxes indicate a motif that is significantly overrepresented (hypergeometric test).

To complement the promoter motif analysis we also tested clusters for enrichment of TF targets inferred from DNA affinity purification sequencing (DAP-seq) data (O’Malley et al., 2016) (Supplemental Figure S1). In DAP-seq, TF target genes can be inferred based on the locations of TF-binding peaks, and data is available for a total of 349 Arabidopsis TFs in the Plant Cistrome Database (http://neomorph.salk.edu/PlantCistromeDB). The patterns of TF-target enrichment across clusters were in broad agreement with the promoter motif analysis, however, some notable differences were also apparent. This included overrepresentation for targets of some ERF family TFs among several coexpression clusters. ERF DNA-binding motifs were not detected as widely enriched in our promoter analysis, which may reflect additional complexity in TF-DNA binding. These findings highlight the potential added value of utilizing complimentary TF interaction data sets.

Regulation of gene expression often occurs through the coordinated action of multiple TFs. To obtain insight into this multi-factorial transcriptional control of the SA-induced response, we next used the DAP-seq data to determine the overlap in SA-induced target genes by the SA-induced TFs. For the 129 SA-responsive TFs for which DAP-seq DNA binding profile data were available, we computed the Jaccard similarity index (i.e., the fraction of shared target genes) over all possible pairs of target genes. Unsupervised hierarchical clustering largely grouped TFs according to their family, but weaker overlap was observed between the members within the different families (Figure 5). The WRKY family displayed the highest degree of within-family overlap, targeting broadly similar sets of genes. Differences among TFs belonging to the same family were also evident, as shown for the NAC, the ERF, and the bZIP TFs, for clear differences in the degree of similarity between family members was highlighted, suggesting within-family specificity targeting distinct sets of SA response genes.

**Figure 5.**
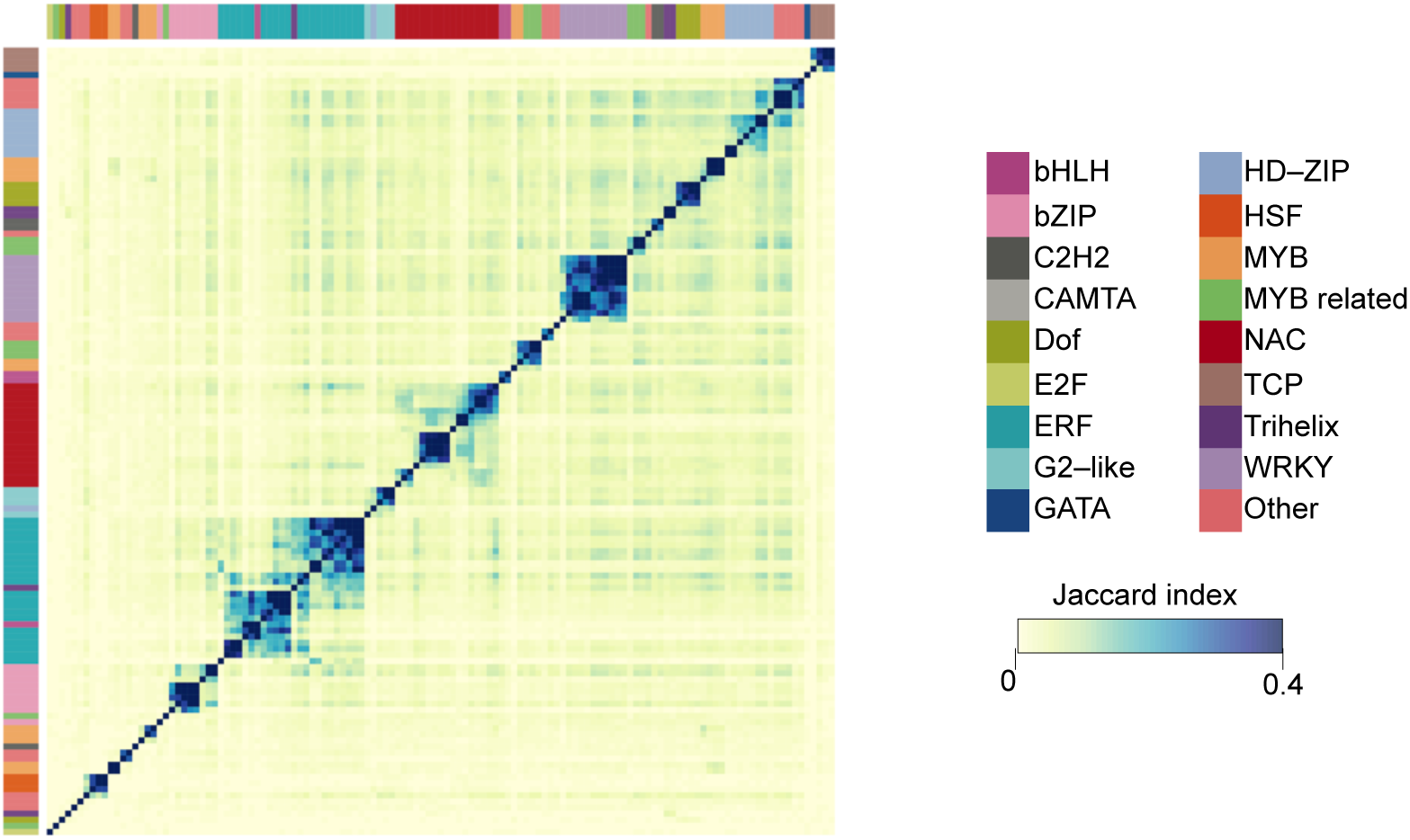
Overlap between the target genes of SA-responsive TFs. Heatmap depicting the overlap in TF target genes by different members of TF families (inferred from DAP-seq data) as measured by the Jaccard similarity index, where both TF and target genes were differentially expressed in response to SA treatment. The pairwise matrix of Jaccard indices are organized via unsupervised hierarchical clustering. Sidebars indicate TF family membership.

In summary, our analyses suggest a complex array of different *cis-*regulatory motifs, their combinations and cognate TFs that allow SA to coordinate the expression of distinct gene modules that comprise the SA response.

### A temporal model of SA-Mediated Transcriptional Reprogramming

The above analyses clearly showed that the SA response is associated with distinct transcriptional signatures, which can then be linked to specific mechanisms of regulation. We next sought to combine the SA-regulated expression profiles with TF-DNA binding data to generate a unified temporal model of the underlying SA-controlled GRN. To achieve this, we used Dynamic Regulatory Events Miner (DREM) (Schulz et al., 2012), a powerful network-reconstruction tool, which integrates static protein–DNA interaction with time course expression data by searching the time series for bifurcation events and predicting the TFs that regulate these paths. Figure 6 shows the DREM-reconstructed temporal transcriptional network, built with the 129 SA-responsive TFs where interaction information could be inferred from DAP-seq data (O’Malley et al., 2016).

**Figure 6.**
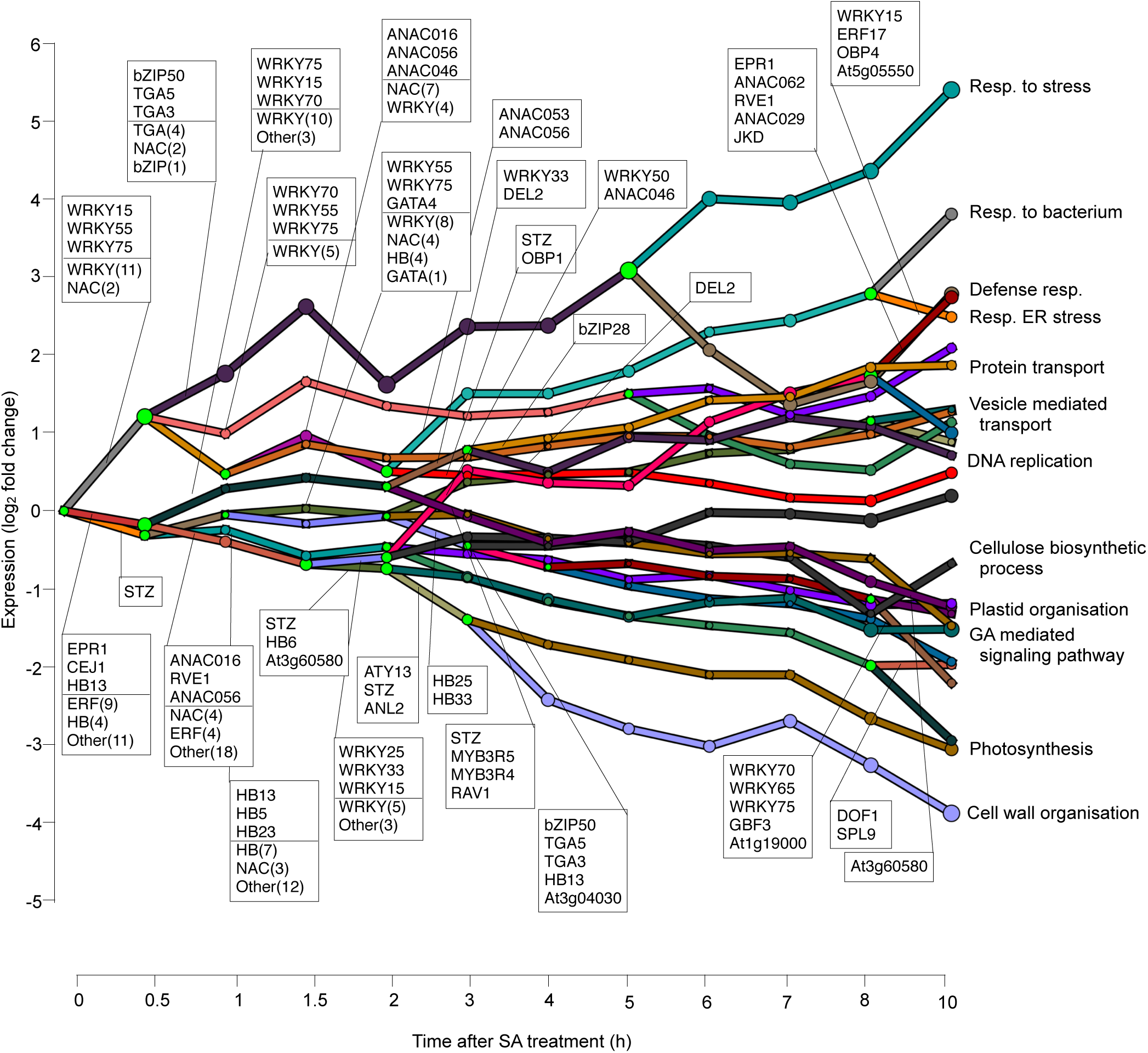
Modeling the SA pathway GRN. Simplified DREM model annotated with TFs based on DAP-seq data. Each path corresponds to a set of genes that were coexpressed. Green nodes represent a bifurcation point where coexpressed genes diverge in expression. Only the top three TFs (strongest associations) and summary of TF families involved are shown. In order to improve readability, only time points until 10 h are shown as no additional path splits were identified after this time. Selected paths are annotated with the most significant functional category overrepresented among the genes assigned to a given path.

This SA GRN model contains 27 unique paths controlled by a wide range of TF families. Bifurcation points were identified at several time points and suggest complexity in the network. In line with our analysis of time of first differential expression (Figure 1), the majority of transcriptional switch points occur early on in the time course, within 4 h of SA treatment. The biological coherence of the identified paths was confirmed with GO term analysis, which revealed that most paths were enriched for distinct biological processes, with many of these being SA related and consistent with the SplineCluster analysis. Among the upregulated paths this included broad terms like ‘response to biotic stimulus’ and ‘response to stress’, as well more specific ones such as ‘vesicle mediated transport’ and ‘DNA replication’, whereas downregulated paths included ‘photosynthesis’ and ‘cell wall organization’.

Regulation of the first path upregulated by SA is enriched for WRKY TFs. Among these is the well-studied WRKY70, which plays an important role as a positive regulator of SA-mediated defense, but is also a negative regulator of SA accumulation (Li et al., 2004; Wang et al., 2006; Knoth et al., 2007). The model also predicted WRKY75 as a regulator of the immediate transcriptional response to SA. This is in line with the recent finding that WRKY75 positively regulates SA accumulation through direct activation of *ICS1* (Guo et al., 2017), but our model also predicts a broader role of WRKY75 in the immediate transcriptional regulation of genes induced following SA pathway stimulation. Two NAC TFs, ANAC075 and ANAC002/ATAF1 are also assigned to this first path of upregulation. Thus, our model suggests that WRKY and NAC TFs are the dominant regulators of genes that are induced in the first stage of the SA response. Interestingly, when this first upregulated path splits at 0.5 h, the WRKY and NAC TF families were assigned to different paths in the bifurcation point, suggesting specificity in regulatory roles at this time point. bZIP family-member TGA TFs also appeared on the same path as the NAC TFs, suggesting possible connective regulation of genes in this group. As time progresses, additional bifurcation points in the model give rise to an increasing number of paths, which are then annotated with distinct sets of TF members from different families, including WRKY, NAC, E2F and ERF. Downregulated gene paths were also annotated with coherent sets of TFs. For example, several early downregulated paths were annotated with homeobox (HB) TFs, and several of these TFs were themselves downregulated by SA treatment. Another notable prediction was the coordinated early downregulation of SA target genes by the SA-upregulated zinc finger TF STZ, which has been shown to function as a repressor of transcription (Saibo et al., 2009)

Thus, while static binding data are available for only a subset of SA-inducible Arabidopsis TFs, the dynamic network model that was inferred by DREM identified known regulators of the SA response and in addition, predicted novel ones, as well as being able to differentiate between early and secondary response regulators of the SA GRN.

### The Dynamic SA/JA Crosstalk Transcriptome

To explore the crosstalk between the JA and SA signaling pathways at the transcriptional level, we generated RNA-seq libraries from Arabidopsis leaves treated with a combination of MeJA and SA. These samples were harvested in parallel with the mock- and MeJA-treated samples described in Hickman *et al.,* (2017) and the SA-treated samples described in this study. As expected, simultaneous induction of both JA- and SA-pathways could result in altered transcriptional response dynamics when compared to the single hormone treatments alone. Figure 7A shows an example of a gene that is affected by SA/JA crosstalk. Transcript levels of the established JA-pathway marker gene, *VSP2,* are strongly upregulated by MeJA treatment alone, while in the MeJA + SA combined treatment initially follow a highly similar pattern of upregulation. However, approximately two hours after treatment with MeJA + SA, suppression of *VSP2* is clearly evident and transcript levels were lower than the MeJA single-treatment profile for the remainder of the time course.

**Figure 7.**
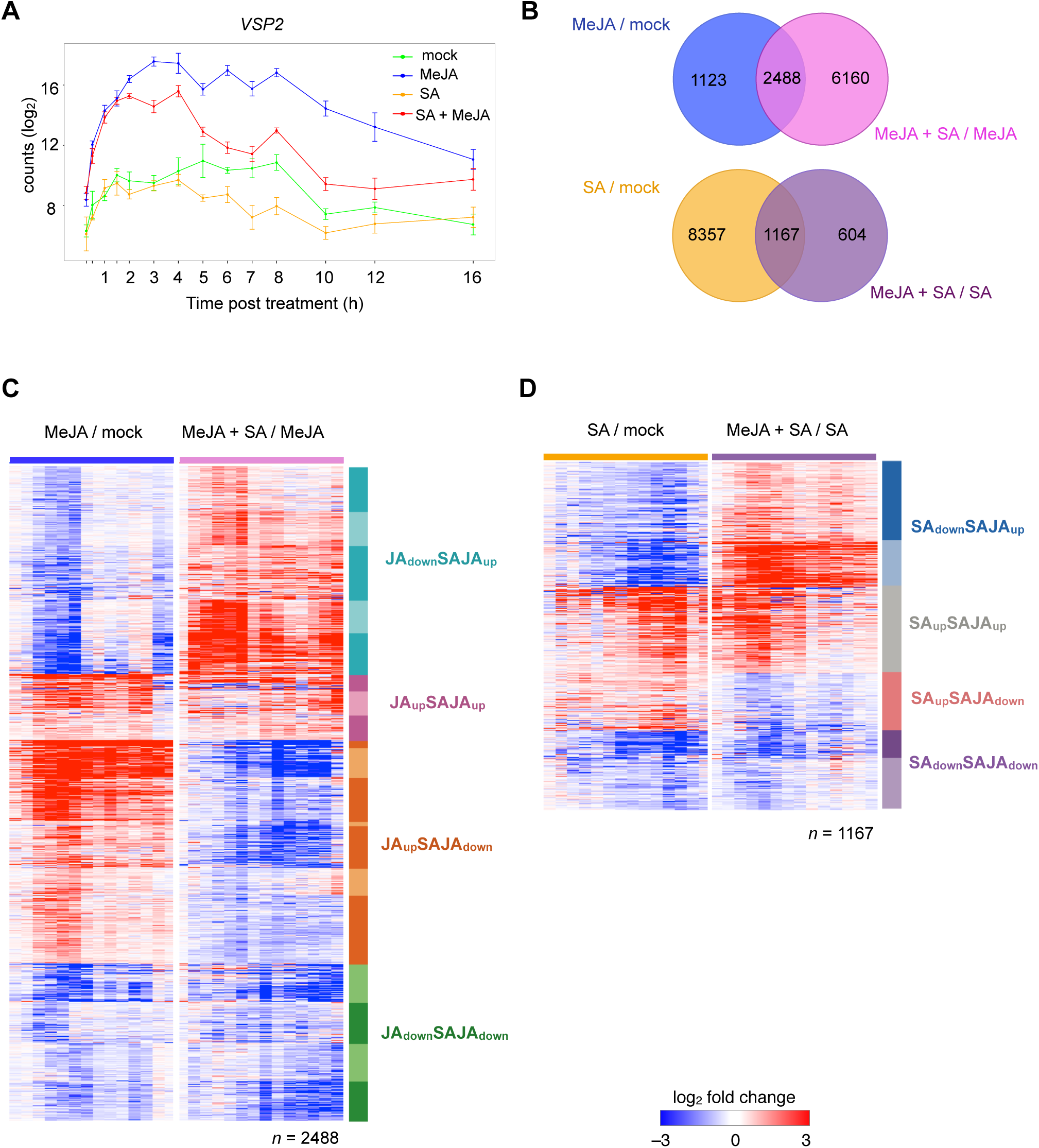
Temporal analysis of SA/JA pathway crosstalk at the transcriptional level. **(A)** Example expression profile of a gene that is subject to SA/JA crosstalk. *VSP2* is upregulated by MeJA treatment (blue) when compared to mock treatment (green). When a combined treatment of MeJA and SA is applied (red), *VSP2* is first upregulated and then subject to SA-mediated suppression (crosstalk). SA alone (orange) represses *VSP2* expression compared to mock levels. y-axis, transcript abundance; x-axis, time (h) post application of hormones; error bars indicate SE. **(B)** Venn diagrams showing the number and overlap between single and combined hormone treatments. Top: genes found differentially expressed in the comparison of MeJA-versus mock-treated plants (blue) and MeJA versus combined SA- and MeJA-treated plants (pink). Bottom: genes found differentially expressed in the comparison of SA-versus mock-treated plants (orange) and SA-versus combined SA- and MeJA-treated plants (purple). **(C)** Multi-condition clustering of 2488 genes found differentially expressed in the MeJA-versus mock and MeJA versus SA+MeJA comparisons. Genes were partitioned into distinct coexpressed gene clusters following clustering of gene expression profiles responding to MeJA alone and to the combined SA- and MeJA treatment. **(D)** Multi-condition clustering of 1167 genes found differentially expressed in the SA-versus mock and SA versus SA+MeJA comparisons. Genes were partitioned into distinct coexpressed gene clusters following clustering of gene expression profiles responding to SA alone and to the combined SA- and MeJA treatment. For both **(C)** and **(D)**, individual clusters are grouped into broader response categories of DEGs that capture the core crosstalk profiles and are indicated by different colors (different color shades indicate the original finer-grain clusters). Each row of a heat map represents an individual gene and indicates fold-change in gene expression. Red and blue coloring indicate induction and suppression of expression respectively.

To assess the global extent of the crosstalk between MeJA and SA pathways we identified genes that were significantly up- or downregulated by SA + MeJA, compared with either MeJA or SA alone using the GLM-based approach described above, and assessed the degree of overlap existing between these sets of genes and the genes that respond to the related single-treatments (Figure 7B). Strikingly, 69% (2488/3611) of MeJA-responsive genes were significantly affected by SA/JA crosstalk, whereas the effect on the SA-controlled transcriptome was less severe, albeit it significant, with 12% (1167/9524) of SA-responsive genes significantly affected by the combined treatment. Thus, in our timeseries study, the MeJA-regulated gene expression program was dramatically altered by the addition of SA, whereas SA-regulated transcriptional reprogramming is to a lesser extent perturbed by the addition of MeJA.

To obtain detailed insight into the dynamic interplay between SA and JA we used SplineCluster in multi-condition clustering mode to identify crosstalk-sensitive MeJA- and SA-responsive genes that are coexpressed across the single and combined treatment timeseries (Figure 7C-D). Clustering was performed using the single-treatment expression profile (condition 1) and the expression in the combined treatment relative to the single treatment (condition 2). Thus, the clustering algorithm will group genes based on their behaviour following activation of the single hormone pathway and how this pattern of up- or downregulation is perturbed with the addition of the other hormone. The clustering identified distinct sets of coexpressed genes (fine-grained crosstalk clusters), which we then grouped into broader response categories reflecting either the enhancement or suppression of single-treatment responsive genes (coarse-grained crosstalk clusters), and subsequently analysed the clusters for enrichment of functional annotations (Supplemental Data Set S3 and S4).

The effects of SA on MeJA-responsive gene expression were broad, with numerous instances of expression antagonism or enhancement for genes that were up- or downregulated by the MeJA single treatment (Figure 7C). In line with previous studies, the vast majority of genes that are induced by MeJA were sensitive to SA-mediated suppression (JA_up_SAJA_down_), and were associated with known JA-inducible genes (defined by gene ontologies). SA also affects the expression of genes suppressed following MeJA treatment, with both enhancement (JA_down_SAJA_up_) and synergistic suppression (JA_down_SAJA_down_) of MeJA-repressed genes evident. Genes in JA_down_SAJA_up_ were enriched for SA-pathway genes and those related to plant immunity, indicating that SA can counteract MeJA-dependent suppression of SA-associated genes. Genes in JA_down_SAJA_down_ are genes that are further repressed by SA in comparison to the downregulated pattern after MeJA alone. Genes in this group were enriched for functional categories associated with development and genes associated with auxin signalling, suggesting that MeJA and SA-signalling can synergistically repress a range of development-related genes.

A smaller subset of JA-induced genes displayed enhanced expression following the double treatment (JA_up_SAJA_up_). GO analysis revealed that this group of genes was enriched for both known JA- and SA-associated genes, suggesting that these genes may represent a convergence point for the two pathways (Supplemental Data Set S3). Interestingly, genes in this group were also overrepresented for genes associated with ET and ABA signalling, suggesting that these genes may be responsive to a broad range of hormone-inducing environmental stimuli.

The effects of MeJA on SA-responsive gene expression were also varied with both antagonistic and synergistic regulation of genes that respond to the SA single treatment (Figure 7D). Genes that are repressed by SA and upregulated by the combined treatment (SA_down_SAJA_up_) were enriched for JA-associated genes, suggesting that MeJA restrains the suppressive effect SA has on the expression of these genes (Supplemental Data Set S4). SA-induced genes that were synergistically enhanced by MeJA (SA_up_SAJA_up_) were enriched for JA- and SA-pathway genes (Supplemental Data Set S4). Interestingly, genes encoding TFs were enriched among the synergistically enhanced set of genes with further analysis showing specific enrichment for the WRKY and MYB TF families among this set. This finding suggests that members of these families may play important roles in the synergistic activation of target genes by both SA and MeJA. SA inducible genes that are sensitive to MeJA-mediated suppression were also evident (SA_up_SAJA_down_), with genes in this group enriched for terms associated with cell communication and signal transduction, including the several immune receptors. This suggests that MeJA may specifically suppress signalling proteins associated with the SA pathway. Finally, SA-responsive genes that are synergistically downregulated by MeJA are overrepresented for a broad range of processes generally related to development (including auxin) and responses to light.

Next, we analysed the promoters of genes assigned to either the fine- or course-grained crosstalk clusters to identify TF-motifs with putative roles in SA/JA crosstalk (Supplemental Figures S3 and S4). Specific motif enrichment was observed in crosstalk clusters at both levels of granularity. In general, MeJA-induced genes that were sensitive to SA-mediated suppression (JA_up_SAJA_down_) were enriched for bHLH family binding motifs, while SA-induced genes sensitive to MeJA-mediated suppression (SA_up_SAJA_down_) were enriched for WRKY family binding motifs. This raises the possibility that antagonistic SA/JA pathway crosstalk involves the targeting of these *cis*-regulatory elements or the TFs that bind them. Interestingly, the promoters of genes in JA_up_SAJA_up_ were enriched for binding motifs for both WRKY and bHLH TFs, while JA_down_SAJA_up_ and JA_up_SAJA_down_, showed enrichment for only WRKY or bHLH binding motifs, respectively. This observation suggests that MeJA-upregulated genes that are synergistically enhanced by SA are targeted by both WRKY and bHLH TFs as part of a joint activation programme following the accumulation of both hormones.

### Network Analysis Identifies Novel Regulators of the SA Response

Our analysis of SA-mediated transcriptional reprogramming uncovered known aspects of the SA gene regulatory network and predicted many more. For example, our coexpression clustering analysis identified coherent gene modules, which can be used to associate genes of unknown function with particular biological processes. This guilt-by-association principle is an often used approach that we, and others, have previously demonstrated as a successful way to uncover novel regulators of particular biological processes and pathways in Arabidopsis (Lewis et al., 2013; Song et al., 2016; Hickman et al., 2017; Walker et al., 2017). To identify novel regulators of the SA response, we selected TF-encoding genes with no previously reported roles in the SA response pathway that were present in clusters that are enriched for terms related to defense responses. We paid particular attention to members of the NAC and WRKY TFs because these families were most significantly overrepresented in the SA-induced DEG set (Figure 2). In total we identified homozygous Arabidopsis T-DNA insertion lines for 18 thus far uncharacterized TF-encoding genes. In some cases, pairs of candidate TFs displayed high sequence similarity and we predicted that single mutants might not display the full effects on host immunity. To account for this possibility, we identified three pairs of genetically-unlinked paralogous genes from our candidate list (Bolle et al., 2013) and either generated double mutants by crossing the single mutants or obtained the double mutants from the GABI-DUPLO collection (Bolle et al., 2013).

We then functionally analyzed the selected single and double mutants for their resistance against the hemi-biotrophic bacterial pathogen *Pseudomonas syringae* pv. *tomato* DC3000 (*Pst* DC3000) that is controlled by SA-inducible defenses (Pieterse et al., 2012). A list of the 21 mutants examined for altered susceptibility to *Pst* is shown in Supplemental Table S1. Of the 21 TF mutants tested, the single mutant of the TF *ANAC090*, and the double mutant of *ANAC090* and its paralog *ANAC061*, had an altered level of resistance to *Pst* DC3000, being more resistant.

The expression kinetics of the *ANAC061* and *ANAC090* genes in wild-type Col-0 plants in response to SA was similar and both genes were therefore assigned to SA response cluster c22 (Figure 8C), which is enriched for genes associated with plant defense (see analysis of the SA coexpression clusters described above). Furthermore, a public database search using the Arabidopsis eFP Browser (Winter et al., 2007) revealed that both *ANAC061* and *ANAC090* were induced by a range of biotic stresses, and particularly highly induced upon treatment with bacterial-derived elicitors, flg22 or HrpZ (assayed 1 h post treatment), further supporting a role for these TFs in the immune response. The expression behavior of the *ANAC61* and *ANAC90* genes in the *anac061*, *anac090*, and *anac061 anac090* mutant lines was verified by qRT-PCR analysis (Supplemental Figure S4). This showed extremely low or undetectable expression levels relative to wild-type following immune elicitation by flg22, if the (single and double) mutants carried the mutation in that gene, and overcompensation in expression of the homologous gene in the two single mutants.

**Figure 8.**
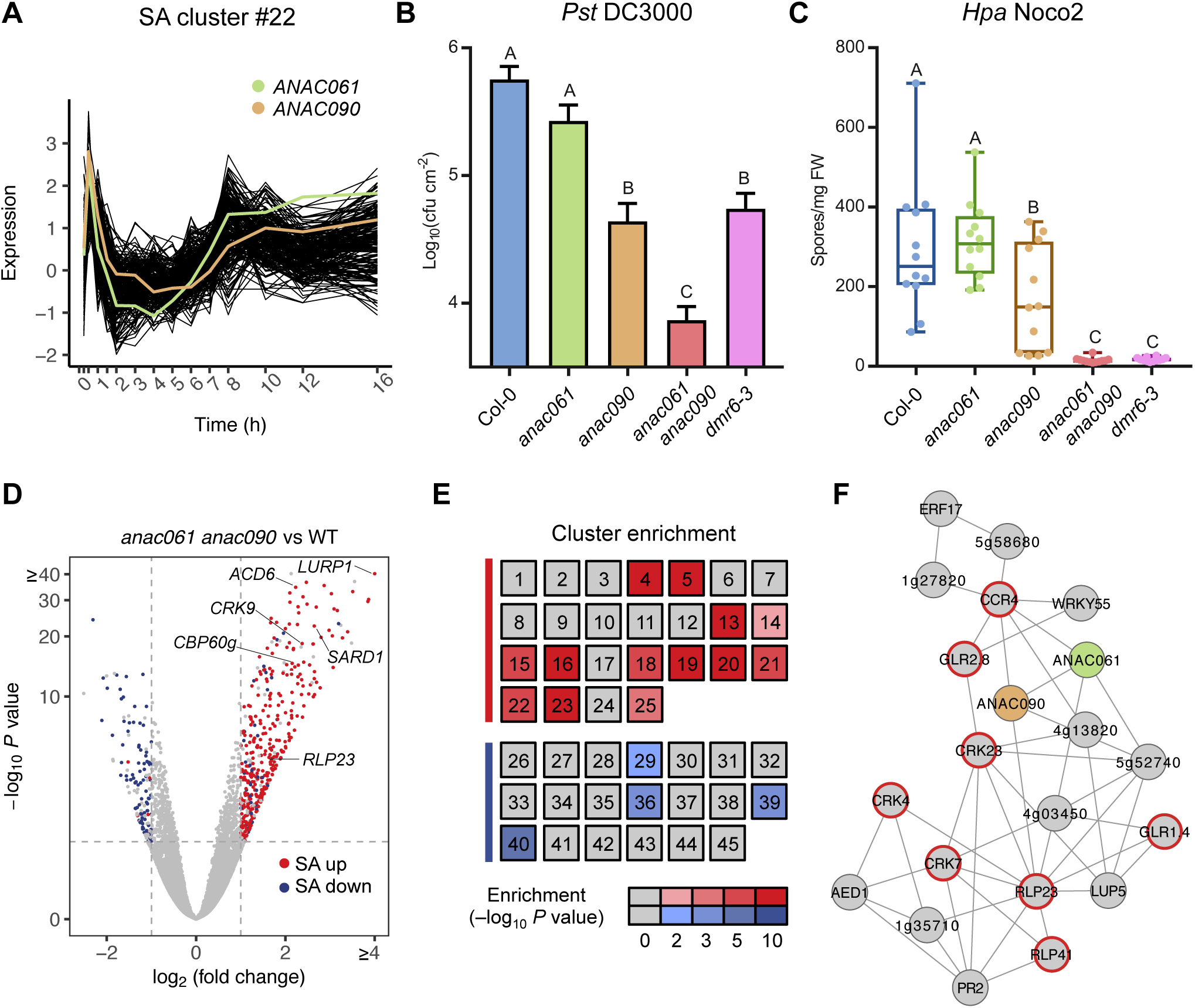
Identification of ANAC061 and ANAC090 as coexpressed homologous TFs that regulate resistance to *P. syringae* and *H. arabidopsidis*. **(A)** *ANAC061* and *ANAC090* belong to the same coexpression cluster (c22) following SA treatment. **(B)** Quantification of bacterial growth in leaves of Col-0 (WT), *anac61, anac90, anac61 anac90*, and *dmr6-3* lines at 72 h after pressure infiltration with *P. syringae* pv. *tomato* (*Pst*) DC3000. One-way ANOVA and post-hoc Tukey test was used to compare the log_10_-transformed *Pst* growth data between the different genotypes (*n*=8; error bars indicate SE). **(C)** Amount of *H. arabidopsidis* Noco2 spores on Col-0 (WT), *anac61, anac90, anac61 anac90*, and *dmr6-3* lines at 7 dpi. One-way ANOVA and post-hoc Tukey test was used to compare the count data of different genotypes to WT (*n*=8; error bars indicate SE). **(D)** Volcano plot depicting differentially expressed genes (FDR < 0.05, |log_2_ fold change| > 1; indicated by dashed grey lines) in *anac61 anac90* compared to Col-0 (WT) in basal conditions. Genes that are up- and downregulated following SA treatment of Col-0 (WT) are coloured red and blue, respectively. Selected genes with defence-related functions are labeled. **(E)** Heatmap indicating hypergeometric enrichment *P* value (after Bonferroni correction) of genes with significantly increased (red gradient) or decreased (blue gradient) transcript levels in *anac61 anac90* in each SA-responsive coexpression cluster (see Figure 2A). Clusters are grouped according to classification as upregulated (red bar) or downregulated (blue bar) following SA treatment. **(F)** Coexpression network obtained using public whole-genome transcriptome data sets with *ANAC061* and *ANAC090* as baits. Genes encoding receptor or receptor-like proteins are marked with a red outline.

Figure 8B shows that the single *anac090* mutant displayed a reduction in the growth of *Pst* DC3000, while there was no difference with wild-type Col-0 plants in the *anac061* mutant. The *anac061 anac090* double mutant accomplished an even greater reduction of bacterial growth than the single *anac090* mutant, and was less diseased than the positive control mutant *dmr6-3* (Zeilmaker et al., 2015). We then also tested the *anac061*, *anac090,* and *anac061 anac090* mutants for their resistance against the SA-controlled oomycete pathogen *Hyaloperonospora arabidopsis* (*Hpa*) isolate Noco2. As shown in Figure 8C, the level of susceptibility to *Hpa* Noco2, based on quantification of sporulation, was reduced in the *anac090* mutant, whereas *anac061* developed symptoms similar to wild-type plants. Again, the *anac061 anac090* double mutant displayed even greater levels of resistance, reaching a similar reduction in *Hpa* sporulation as the *dmr6-3* mutant. Taken together, these results suggest that *anac061* and *anac090* have shared activities as negative regulators of immunity against biotrophic and hemi-biotrophic pathogens.

To provide insight into the biological processes contributing to reduced pathogen performance on *anac061 anac090*, we performed RNA-seq analysis on leaves harvested from the double mutant and wild-type Col-0. A total of 592 genes were differentially expressed between *anac061 anac090* and wild-type plants (453 were upregulated and 139 were downregulated in the double mutant; Supplemental Data Set S5). Functional category analysis showed that for genes with upregulated expression in the double mutant versus wild type, processes such as “defense response” and “response to SA” were among the most significantly enriched terms, while downregulated genes were associated with primary metabolism (Supplemental Data Set S5). Volcano plot visualization demonstrated that the transcriptional signature of *anac061 anac090* was characterised by an upregulation of many genes that were also induced by SA in wild-type plants in our time series data set and include important regulators of SA signalling and immunity (e.g., *SARD1*, *CBP60g*, *ACD6*, *CRK9*, *LURP1*) (Figure 8D). Genes that were downregulated in *anac061 anac090* generally overlap with genes that were downregulated by SA. In accordance with these findings *anac061 anac090*-upregulated DEGs were specifically overrepresented in several coexpression clusters of the SA-induced DEGs, and many of these are enriched for defense-related functions. Furthermore, *anac061 anac090*-downregulated DEGs were only overrepresented among downregulated coexpression clusters (Figure 8E). Thus, these RNA-seq data indicate that ANAC061 and ANAC090 are negative regulators of SA-mediated immunity and specifically target certain sectors with the SA network. This hypothesis was further supported by gene coexpression network analysis performed with *Arabidopsis* transcriptome data using ATTED-II (Obayashi et al., 2018), which demonstrated coregulation of the *ANAC061* and *ANAC090* genes with genes with immune annotations including a large proportion that encode receptor-like proteins (Figure 8F).

## Discussion

Plant immunity is regulated by a complex network of cross-communicating signalling pathways that control gene expression programs responsible for mediating a range of adaptive defence responses. SA is a key immune signalling hormone that controls defence responses that are effective against pathogens with (hemi)-biotrophic lifestyles (Vlot et al., 2009). The SA-response is influenced by the activity of other phythormones, with the interplay with JA-pathway particularly important. SA-JA crosstalk can optimize the immune response against single attackers and provides a potentially powerful mechanism to prioritize one pathway over the other, depending on the context in which they are activated (Pieterse et al., 2012).

Previous transcript profiling studies of the SA response and SA-JA crosstalk in Arabidopsis have analyzed these processes with a limited number of time points after exogenous application of SA or BTH. (Schenk et al., 2003; Wang et al., 2006; Blanco et al., 2009; Van der Does et al., 2013). These studies gave a static picture of the overall changes in gene expression that occur in response to SA and the interaction between the SA and JA pathways, and provided limited information on the timing and sequence of transcriptional changes and related biological processes affected, and lack the resolution required to infer an in depth understanding of the dynamic GRN underlying these responses. To move knowledge beyond these previous studies, this article describes the generation and analysis of information-rich high-resolution time-course gene expression data sets to first analyse the response of Arabidopsis leaves to SA treatment alone, and then to analyse the transcriptional effects of crosstalk between SA and JA by monitoring expression changes in response to a combined SA + MeJA treatment.

### Temporal Transcription Landscape and Dynamic GRN Modeling of the SA Response

Analysis of this gene expression time series showed that SA induces large-scale transcriptional reprogramming with approximately one-third of the Arabidopsis genome changing in expression during the first 16 h following application of SA. Our analysis of differential gene expression greatly expanded the number of genes that respond in an SA-dependent manner, including hundreds of transcripts that could not have been detected previously with microarrays. From these data, we were able to make a series of detailed regulatory insights into the SA response, including the timing of transcriptional response, biological processes targeted, patterns of coregulation, and novel network regulators.

One striking finding from our analysis was that SA-mediated differential expression occurs in clearly defined temporal waves, which were enriched for genes corresponding to distinct biological processes (Figure 1). The majority of gene expression changes (upregulation and downregulation) occurred rapidly within the first 2 h after SA treatment, taking place in the first wave (Figure 1). Within the first wave of gene upregulation were many well-established immune response genes. The second wave of upregulation (2–4 h) was associated with genes involved in the wounding response. Interestingly, within the third wave (4 h onwards) were many genes associated with cell cycle and DNA repair. The two discernable waves of downregulated genes were both associated with different categories of metabolism: photosynthesis and development (wave 1) and starch (wave 2). The cluster analysis identified groups of SA-responsive genes that had not been observed previously with more limited time series data (Figure 2). Analysis of the individual clusters identified groups of genes involved in a common process among both upregulated and downregulated genes. Annotations among upregulated clusters included classical immune responses (e.g., signal transduction, protein phosphorylation, etc), as well more specific terms, such as unfolded protein response and DNA repair; downregulated clusters were associated with processes related to primary metabolism and development, such as photosynthesis, chlorophyll metabolism, and auxin signalling. The ability to see these differences in transcriptional activity over time provides a highly nuanced view of the genes and respective biological processes involved in the SA response.

It seems likely that genes involved in the same process, with similar expression profiles, are coregulated rather than simply coexpressed during the SA response, and this prediction is strengthened by the analysis of promoter motifs and TF binding. In general, we find significant correspondence among expression pattern, gene functions, and promoter enrichment for known TF motifs in and TF binding (DAP-seq data) (Figure 4 and S1). This combined analysis re-identified many known regulatory components and suggested additional roles for other *trans*- and *cis*-acting factors in regulation of the SA response.

Previous studies have highlighted several members of the WRKY TF family as components in a SA-mediated regulatory network that function as either positive or negative regulators of target genes (Wang et al., 2006). The importance of WRKY TF families in promoting the SA-response in Arabidopsis was clearly illustrated through the specific and highly significant enrichment for WRKY binding sites for these regulators in the promoters of multiple clusters of upregulated genes (Figure 4). This hypothesis was strengthened by the topology of the DREM-derived temporal SA GRN (Figure 6), which indicates that several different WRKY TFs act as master regulators of the SA-pathway. The DREM model is valuable because it makes specific predictions about the TFs that act as regulatory nodes within the GRN. For example, WRKY15, WRKY75 and WRKY55 were among the top ranked TFs assigned to the first path of gene upregulation, suggesting important roles in regulating the immediate/early SA-response. Validating their importance to the SA-response and SA-mediated immunity through, for example, the use of mutant and transgenic TF lines will be important focus of future research.

The identification of many different groups of tightly coexpressed genes with diverse expression patterns during the SA-response, which were enriched in particular groups of functional terms, indicated that a varied set of TFs may specifically regulate the genes in each group. Upregulated gene clusters that were not enriched for WRKY binding motifs were often overrepresented for motifs corresponding to other TF families, such as NAC, CAMTA, HSF, and E2F (Figure 4), highlighting the diversity of TFs and their respective binding elements that regulate SA-mediated transcriptional reprogramming. The DREM-derived GRN could also support a model whereby a variety of TF families act to fine-tune the expression of dedicated sets of target genes in specific sectors of the SA GRN. Following the initial WRKY-dominated activation response, the temporal network model is subject to a series of splits, which represent divergence of genes that were co-regulated up until that point (Figure 6). Many of these expression paths are enriched for specific biological processes, and this set shares a high level of overlap with the different processes also found in SplineCluster analysis. While some WRKY TFs are predicted to regulate splits in gene expression trajectories, the model generally predicts a different set of secondary regulators to be responsible for determining the patterns of genes linked to specific pathways and processes, and include members of the bZIP, NAC and E2F families.

An important caveat is that the DREM analysis was limited to the 349 Arabidopsis TFs for which high-quality binding profiles are currently in the DAP-seq database (O’Malley et al., 2016). Thus, hundreds of known and potentially important regulators of the SA pathway are missing from our analysis. The generation of genome-wide binding profiles for additional SA-regulated TFs will be required to identify the full complement of master and secondary TFs that orchestrate the SA response.

Obviously, transcriptional regulation is only one of several mechanisms of gene regulation in plants, and complex patterns of gene expression can also be controlled at posttranscriptional or posttranslational level. For example, the initial steps in the SA response are regulated through the posttranslational modifications of NPR1, NPR3 and NPR4 (Tada et al., 2008; Fu et al., 2012; Wu et al., 2012; Ding et al., 2018). A future challenge is to link non-transcriptional regulation such as mRNA stability, mRNA translation, and protein stability to transcriptional network models.

### Emerging role for NAC TFs in the SA response

The plant-specific NAC family of TFs play diverse roles regulating plant growth, development and responses to environmental stress. In recent years, the majority of studies have focused on the roles of NAC family members in the regulation of development processes such as cellular differentiation and senescence, and abiotic stresses such as drought and high-salinity (Olsen et al., 2005; Nakashima et al., 2012; Kim et al., 2016). In recent years, several NAC TFs have also been identified as important components of the SA-pathway; for example, the three homologs, ANAC019, ANAC055, ANAC072 function as negative regulators of the SA pathway, while ANAC032 is a positive regulator of the SA response. (Zheng et al., 2012; Allu et al., 2016). Several lines of evidence in this study pointed to the NAC TF family playing an extended role in regulating the SA response. First, our comprehensive cataloguing of SA-responsive genes revealed that, after the WRKY TF family, genes encoding NAC TFs displayed the most significant enrichment among those upregulated in response to SA (Figure 3A). Secondly, motif analysis of promoters of coexpressed genes identified NAC TF motifs as enriched within several clusters (Figure 3B and 4). Third, our dynamic regulatory model of the SA-response predicted several NAC TFs were responsible for regulating several dynamic paths of coexpression (Figure 6).

Using a guilt-by-association approach, we identified several NAC TFs as candidate regulators of the SA-response and screened their respective mutants, and in some cases, double mutants for altered resistance against SA-inducing pathogens. This reverse-genetic screen identified the single *anac090* mutant, and the double *anac061 anac090* mutant as displaying enhanced resistance to *Pst* DC3000 and *Hpa* Noco2 when compared to wild-type plants (Figure 8A-B).

The *anac061 anac090* double mutant was the most resistant, indicating that this paralogous TF pair acts redundantly, and likely regulate overlapping sets of SA-responsive genes. Transcriptome analysis of the double mutant confirmed upregulation of many SA-pathway genes (Figure 8D-E), which likely explains the enhanced resistance against (hemi)-biotrophic pathogens. In support of our findings, a recent study of a NAC TF network during developmental leaf senescence by (Kim et al., 2018) also linked ANAC090 to SA signalling. In that study, they identified ANAC090 as a negative regulator of SA-mediated leaf senescence and they showed that loss of ANAC090 led to increased SA levels and accelerated leaf senescence. Furthermore, through ChIP-qPCR experiments, they showed that ANAC090 binds to DNA upstream of several important SA-signaling genes, including *ICS1* and *EDS5*.

High sequence similarity and genetic redundancy between TF family members can often mask the contribution of individual TFs towards regulating a specific process. This may explain why most of other *ANAC* single mutants tested did not display significant effects on host immunity in the above-described analyses, even if these individual TFs constitute valid nodes in the SA GRN. An important step in future research will be to generate higher order mutants of related NAC TFs and others predicted to play a role in regulating the SA response.

### Dynamic SA-JA Pathway Crosstalk at the Transcriptional Level

In this study, we have defined the immediate transcriptional effects of crosstalk between SA and JA, identifying a large set of target genes whose expression is dynamically altered (upregulated or downregulated) with combined SA + MeJA-treatment, compared with either hormone alone (Figure 7). The effect of SA on the MeJA-regulated transcriptome is highly significant, with 69% of MeJA-responsive genes significantly enhanced or repressed in the presence of SA. Previous studies have typically focused on the antagonistic effect of SA on the JA pathway (Van der Does et al., 2013; Caarls et al., 2017), and our temporal transcriptome data showed that the vast majority of MeJA-response genes are subject to SA-mediated suppression (Figure 7B). The effect of MeJA on SA-regulated genes was less striking, with 12% of genes that were differentially expressed in response to the SA single treatment significantly altered when compared with the combined treatment (Figure 7B).

MeJA-induced target genes were rapidly suppressed by SA treatment, with our temporal data indicating that the majority of SA-mediated suppression occurs within 0 – 3 h after stimulation with a combined SA + MeJA treatment. The altered patterns of MeJA-responsive gene expression in the presence of SA were complex (Figure 7C); clustering of crosstalked expression profiles revealed diverse patterns of coregulation (Figure 7D). A similarly complex picture of coregulation was revealed by cluster analysis of the temporal profiles of SA-responsive genes that were crosstalked by MeJA. These observations likely reflect the combinatorial regulation of these genes by both JA and SA pathways.

Analysis of TF-binding motifs within these clusters (Supplemental Figure S2 and S3) did not reveal potential *cis*-regulatory elements that actively discriminate between sets of genes that are upregulated in response to a given single-hormone treatment and subsequently antagonized in the presence of the alternative hormone. Interestingly, however, we found that the promoters of single-treatment upregulated genes that are synergistically upregulated in the combined SA + MeJA treatment were enriched for bHLH and WRKY TF binding motifs. This pattern of enrichment was not generally observed for other sets of crosstalked genes. Members of the bHLH and WRKY TF families constitute important master regulators of the JA and SA-pathways, respectively. Thus, a simple explanation for this observation is that positive regulators of each pathway converge on promoters of such genes to synergistically enhance gene expression.

In conclusion, the temporal transcriptome datasets show that crosstalk between SA and JA-pathways leads to dynamic transcriptional changes that can be antagonistic or synergistic, and highlight the potentially important role the distribution of *cis*-regulatory elements within the promoters of crosstalked genes for defining their response to a combination of both SA + MeJA.

## Methods

### Plant Materials and Growth Conditions

All wild type, mutant, and transgenic *Arabidopsis thaliana* plants used in this study are in the Columbia accession (Col-0) background. The following T-DNA insertion mutants and transgenic lines were obtained from the Nottingham Arabidopsis Stock Centre: *bzip17* (At2g40950; SALK_004048C), *anac017* (At1g34190; SALK_022174C), *anac047* (At3g04070; SALK_066615), *anac053* (At3g10500; SALK_009578), *anac061* (At3g44350; SALK_041446), *anac078* (At5g04410; SALK_040812), *anac082* (At5g09330; GK-282H08), *anac087* (At5g18270; SALK_079821) *anac090* (At5g22380; SALK_011849C), *anac104* (At5g64530; SALK_022552), *myb50* (At1g57560; SALK_035416C) *myb61* (At1g09540; SALK_106556C), *wrky64* (At1g66560; GABI_519C02), *wrky66* (At1g80590; SALK_055084C), *wrky67* (At1g66550; SALK_027849C), and *anac053 anac078* double mutant from the GABI-DUPLO collection (Bolle et al., 2013) (N2103080). The *anac061* and *anac090* mutants were crossed to generate the *anac061 anac090* double mutant.

Plants were grown as described previously (Hickman et al., 2017). In brief, seeds were stratified at 4°C for 48 h prior to sowing on river sand. Two weeks after germination, the seedlings were transferred to 60-mL pots containing a soil:river sand mixture (12:5) that had been autoclaved twice for 1 h. Plants were cultivated in a growth chamber under a 10-h day (75 µmol/m^2^/s^1^) and 14-h night cycle at 21°C and 70% relative humidity. Plants were watered every other day and received modified half-strength Hoagland nutrient solution containing 10 mM Sequestreen (CIBA-GEIGY GmbH, Frankfurt, Germany) once a week.

### RNA-seq experimental setups

For the SA and SA + MeJA time series experiments, 5-week-old Arabidopsis Col-0 plants were treated by dipping the rosette leaves into a solution containing 0.015% (v/v) Silwet L77 (Van Meeuwen Chemicals BV) and either 1 mM SA (Mallinckrodt Baker) or a combination of SA and 0.1 mM MeJA (Serva, Brunschwig Chemie). Because MeJA was added to the solutions from a 1,000-fold concentrated stock in 96% ethanol, solutions without MeJA received a similar volume of 96% ethanol. Subsequently, developmental leaf six was harvested from four individual SA or SA + MeJA-treated plants at each of the following time points post-treatment: 15 min, 30 min and 1, 1.5, 2, 3, 4, 5, 6, 7, 8, 10, 12 and 16 h. The SA and SA + MeJA treated samples were harvested simultaneously with MeJA and mock-treated samples described in a previously published time series analysis of phytohormone responses (Hickman et al., 2017). For the comparison of the *anac061 anac090* double mutant with wild-type Col-0, two mature leaves (number 6 and 7) were harvested per plant from two 5-week-old plants per genotype, resulting in two biological replicates.

### RNA-seq library preparation and sequencing

Total RNA was extracted using the RNeasy Plant mini kit (Qiagen), including a DNase treatment step in accordance with the manufacturer’s instructions. For the time series experiment, RNA-Seq library preparation and sequencing was performed as described previously (Hickman et al., 2017). Libraries were prepared using the Illumina TruSeq mRNA Sample Prep Kit and sequenced on the Illumina HiSeq 2000 platform with read lengths of 50 bases. For the analysis of the *anac061 anac090* double mutant, RNA-Seq libraries were prepared using the Illumina Truseq mRNA Stranded Sample Prep Kit, and sequenced on the Illumina NextSeq 500 platform with read lengths of 75 bases. All raw RNA-Seq read data are deposited in the Sequence Read Archive (http://www.ncbi.nlm.nih.gov) with BioProject ID PRJNA224133 and PRJNA395645.

### RNA-Seq Analysis

Quantification of gene expression from RNA-seq data was performed as described previously (Van Verk et al., 2013; Hickman et al., 2017). Sequencing reads were aligned to the *Arabidopsis* genome (TAIR version 10) using TopHat v2.0.4 (Trapnell et al., 2009), summarized over annotated gene models using HTSeq-count v0.5.3p9 (Anders et al., 2015) and normalized using the DESeq R package (Anders and Huber, 2010).

To identify genes whose transcript levels differed significantly over time between a given pair of hormone treatments we used the approach previously described in Hickman et al. (2017). Briefly, for each gene a negative binomial generalized linear model was fit to the normalized gene counts with treatment and time as covariates. We then used ANOVA to compare the full model to a reduced model without the treatment variable. To adjust for multiple comparisons, we adjusted the resulting *P* values using the Bonferroni method. Genes with an adjusted *P* value < 0.05 and a fold change > 2 at one or more timepoints were called as DEGs. Genes differentially expressed between MeJA- and mock-treated leaves were described previously (Hickman et al., 2017). Genes differentially expressed between Col-0 and *anac061 anac090* (|log2-fold change| >1; false discovery rate < 0.05) were identified using DESeq2 (Love et al., 2014) with default settings.

### Clustering of Gene Expression Profiles

Clustering of DEGs was performed using SplineCluster (Heard et al., 2006). Genes for the SA vs mock comparison were clustered with a prior precision value of 10^-5^ on the basis of SA expression, which had been subject to the Variance Stabilizing Transformation (VST) method provided by DESeq (Love et al., 2014). Joint clustering (coclustering) of genes that are differentially expressed between MeJA versus mock and SA + MeJA versus MeJA was performed using the log_2_- transformed normalized differences in expression within each of the two comparisons, and with a prior precision of 10^-5^. Coclustering of genes that are differentially expressed between SA versus mock and SA + MeJA versus SA was performed using the log_2_-transformed normalized differences in expression within each of the two comparisons, and with a prior precision of 10^-6^. In each instance, clustering was performed using a range of prior precision values to identify the most informative set of clusters, by balancing within-versus between-cluster variation. An additional reallocation function (Heard, 2011) that redistributes cluster outliers into more appropriate clusters was also performed.

### Gene Ontology Analysis

GO enrichment analysis on gene clusters was performed using GO term finder (Boyle et al., 2004) and an *Arabidopsis* gene association file downloaded from ftp.geneontology.org on 10^th^ Feb 2017. To remove generic GO terms, only GO categories at level two and above in the GO hierarchy were included in the GO analysis. Overrepresentation for the GO categories ‘Biological Process’, ‘Molecular Function’ and ‘Cellular Component’ were identified by computing a *P* value using the hypergeometric distribution and false discovery rate for multiple testing (*P* < 0.05). GO analysis of DREM paths was performed using DREM’s in-built GO enrichment method (Schulz et al., 2012), using the gene association file described above.

### TF Family Analysis

Overrepresented TF families within a set of genes were analyzed as described in Hickman *et al*., 2017. TF family annotations were retrieved from PlantTFDB version 3.0 (Jin et al., 2014) and tested for enrichment using the hypergeometric distribution with *P* values corrected for multiple testing with the Bonferroni method.

### Promoter Motif and Transcription Factor Binding Analysis

For the analysis of known Arabidopsis TF DNA-binding motifs we retrieved published position specific weight matrices from CIS-DB version 1.02 (Weirauch et al., 2014) and those described in Franco-Zorrilla et al. (2014). Promoter sequences defined as the 500 bp upstream of the transcription start site (TSS) were retrieved from TAIR (version 10). The occurrence of a motif within a promoter was determined using FIMO (Grant et al., 2011), where a promoter was considered to contain a motif if it had at least one match with a *P* value < 10^-4^. Motif enrichment was assessed using the hypergeometric distribution against the background of all Arabidopsis genes.

TF-gene interactions were inferred from DAP-seq (DNA affinity purification sequencing) experiments, which provide the genome-wide binding profiles of in-vitro-expressed TFs (O’Malley et al., 2016). DAP-seq peaks for 349 Arabidopsis TFs with a FRiP (fraction of reads in peaks) score ≥5% were retrieved from the Plant Cistrome DB (O’Malley et al., 2016). To improve the interpretability of our models we took an approach based on that outlined in Narsai et al. (2017), and reduced the size of the TF-gene interaction dataset by keeping the strongest 25% DAP-seq peaks for each TF and used ChIPpeakAnno to assign their target genes using a TSS distance threshold of -2000 with all other parameters kept as default (Zhu et al., 2010). The enrichment of TF targets within coexpressed gene clusters was assessed using the hypergeometric distribution as described above.

### DREM Analysis

DREM was run using two input data sets: The first was a matrix of moderated log_2_-fold changes of SA relative to mock expression per time point using count data that had been transformed using the VST method (Love et al., 2014). The second was a TF-gene interaction file derived from DAP-seq data as described above that only considered TFs that were DE in response to SA treatment. The first measured time point at 15 min post SA treatment was omitted from the final model build, as we found that DREM did not generate paths of up- or downregulation at this time point when included.

### Coexpression network analysis

The coexpression network was obtained using the ATTED-II Network Drawer tool with the default provided RNA-seq datasets (http://atted.jp/cgi-bin/NetworkDrawer.cgi) (Obayashi et al., 2018) and using *ANAC061* and *ANAC090* as query genes.

### Disease bioassays

*Pseudomonas syringae* pv. *tomato* DC3000 (*Pst* DC3000) was cultured in King’s B medium supplemented with 50 mg/L rifampicine at 28 °C overnight. Bacteria were collected by centrifugation for 10 min at 4000 rpm, and re-suspended in 10 mM MgSO_4_. The suspension was adjusted to OD_600_=0.002 and pressure infiltrated into three mature leaves of 5-week-old plants with a needleless syringe as described previously (Van Wees et al., 2013). After three days, leaf discs from infected leaves were harvested from two inoculated leaves per plant to give a single biological replicate. Eight biological replicates were harvested for each genotype. Subsequently, 500 µl of 10 mM MgSO_4_ was added to the samples, after which they were macerated using a TissueLyser (Qiagen). Serial ten-fold dilutions were made in 10 mM MgSO_4_, and 30 µl aliquots plated onto KB agar plates containing 50 mg/mL rifampicine. After 48 h incubation at 28°C, bacterial colonies were counted. Bacterial growth in the mutant lines was compared with that of the wild type (Col-0) using ANOVA followed by Dunnett’s multiple comparison tests for mutant screening and ANOVA followed by Tukey’s multiple comparison test for specific ANAC assays, as indicated in the figure legends.

*Hyaloperonospora arabidopsidis* isolate Noco2 (Hpa Noco2) spores were harvested from infected (*eds2* mutant) plants, eluted through Miracloth, and diluted in water to 50 spores/µL. For the disease bioassay, five-week-old plants were spray inoculated with this spore suspension. Plants were subsequently placed at 100% RH, under short day conditions (9 hrs light/15 hrs dark) at 16 °C. After 9 days the spores from eight individual plants were harvested in 5 mL of water and the number of spores per milligram of plant tissue (fresh weight of aerial parts) was counted using a light microscope. Spore counts in the mutant lines were compared with that of the wild type (Col-0) using ANOVA followed by Tukey’s multiple comparison test

## Supplemental Data

**Supplemental Figure 1.** Enriched TF targets in SA-responsive gene coexpression clusters.

**Supplemental Figure 2.** Enriched *cis*-regulatory motifs in clusters of MeJA-responsive genes that are sensitive to SA crosstalk.

**Supplemental Figure 3.** Enriched *cis*-regulatory motifs in clusters of SA-responsive genes that are sensitive to MeJA crosstalk.

**Supplemental Figure 4.** Expression of *ANAC061* and *ANAC090* in Col-0 (WT), *anac061*, *anac090* and *anac061 anac090*.

**Supplemental Table 1.** A list of all mutant alleles for predicted SA-pathway regulators and their *Pst* DC3000 disease resistance phenotypes.

**Supplemental Data Set 1.** SA-responsive DEGs.

**Supplemental Data Set 2.** Cluster membership for SA-responsive DEGs following clustering using SA-treated gene expression profiles. GO terms overrepresented in the clusters.

**Supplemental Data Set 3.** Cluster membership for MeJA-responsive DEGs that are crosstalked by SA. GO terms overrepresented in the clusters.

**Supplemental Data Set 4.** Cluster membership for SA-responsive DEGs that are crosstalked by MeJA. GO terms overrepresented in the clusters.

**Supplemental Data Set 5.** Genes differentially expressed in *anac061 anac090* compared to Col-0. GO terms overrepresented in the upregulated and downregulated *anac061 anac090* DEG sets.

## Acknowledgements

This work was supported by the Netherlands Organization for Scientific Research through the Dutch Technology Foundation (VIDI 11281 to S.C.M.V.W. and VENI 13682 to R.H.), by the European Research Council (Grant 269072 to C.M.J.P.), and by the CAPES Foundation, Ministry of Education of Brazil (DF 70040-020 to M.P.M.).

