## Supplemental Figures and Tables for "Transcriptional Dynamics of the Salicylic Acid Response and its Interplay with the Jasmonic Acid Pathway"

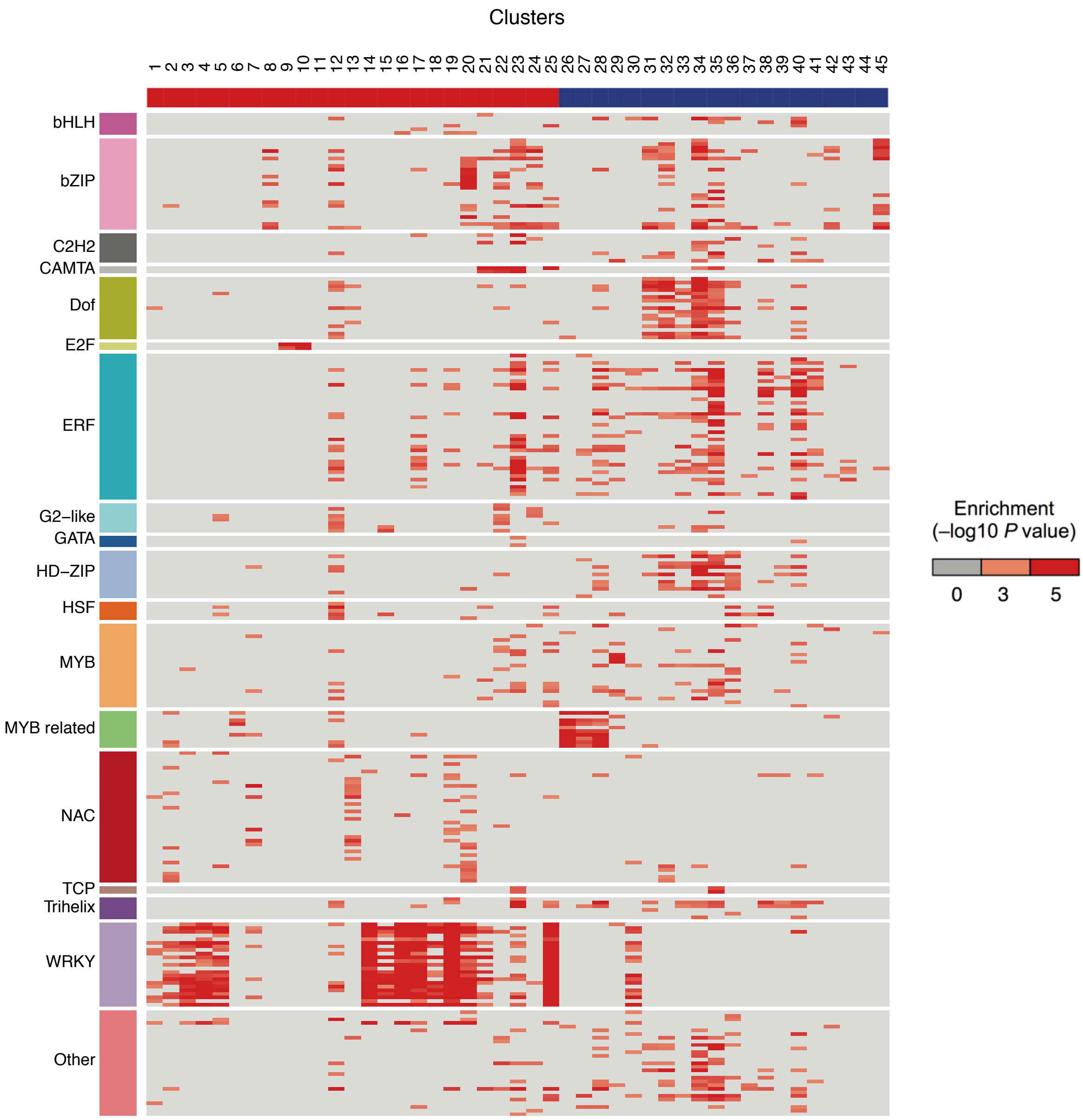


**Supplemental Figure S1. Enriched TF targets in SA-responsive gene coexpression clusters.**
(Supports Figure 4)
TF targets inferred from DAP-seq peak data are differentially enriched in the promoters of genes clustered on the basis of their expression following application of SA. Rows indicate TFs and are colored by corresponding family. Columns indicate coexpression clusters, with red and blue column colors differentiating between up- and downregulated clusters respectively. Red boxes indicate a TF targets are significantly overrepresented among genes in a given cluster (hypergeometric test).

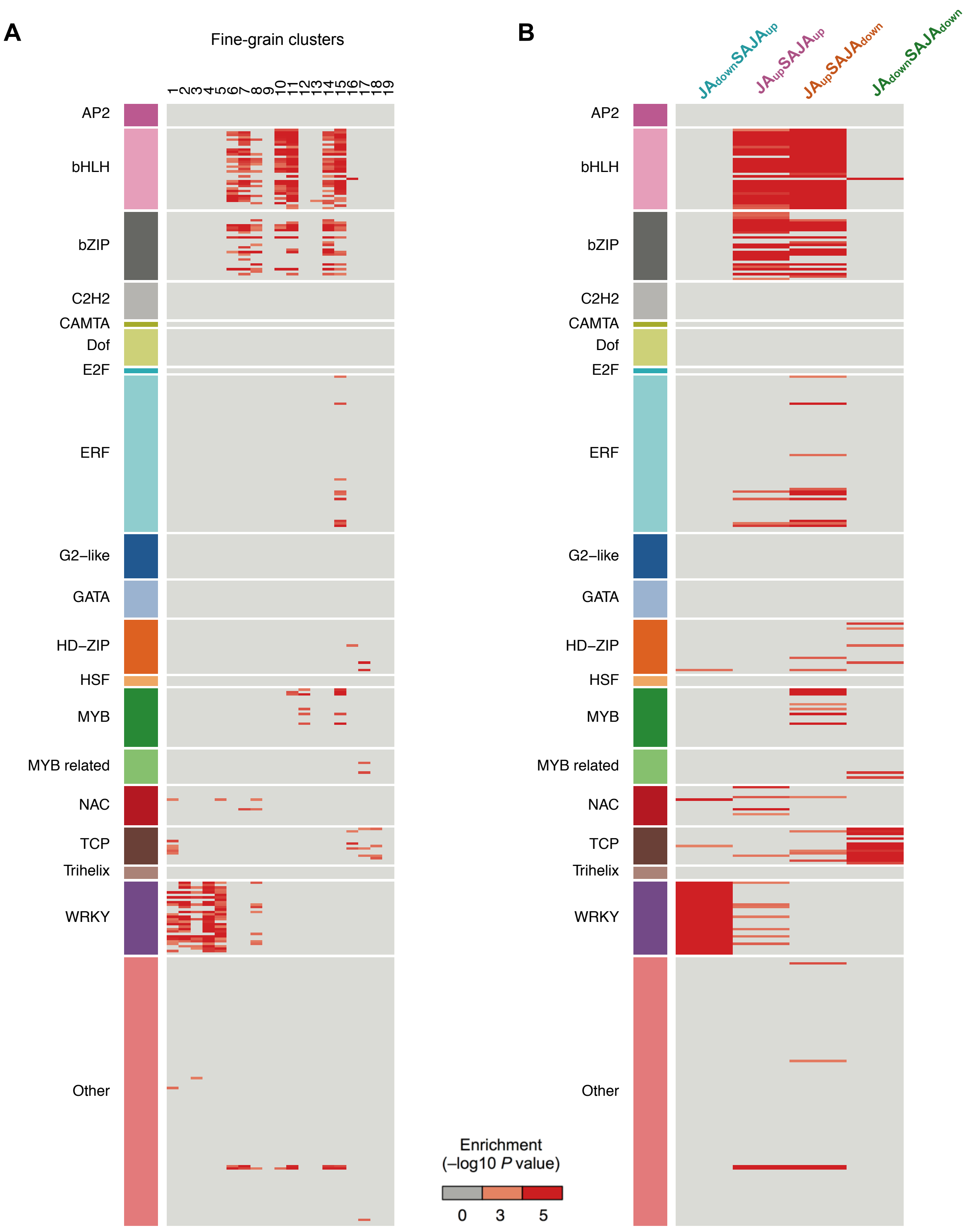


**Supplemental Figure S2. Enriched *cis*-regulatory motifs in clusters of MeJA-responsive genes that are sensitive to SA crosstalk.**
(Supports Figure 7C)

Known TF DNA-binding motifs enriched in the promoters of genes clustered based on their gene expression profiles responding to MeJA alone and to the combined SA- and MeJA treatment as shown in Figure 6C. **(A)** Motif enrichment *P* values in the 19 fine-grain clusters. **(B)** Motif enrichment *P* values in the course-grain clusters. Columns indicate clusters. Rows indicate motifs and are colored by corresponding TF family. Red boxes indicate a motif that is significantly overrepresented (hypergeometric test).

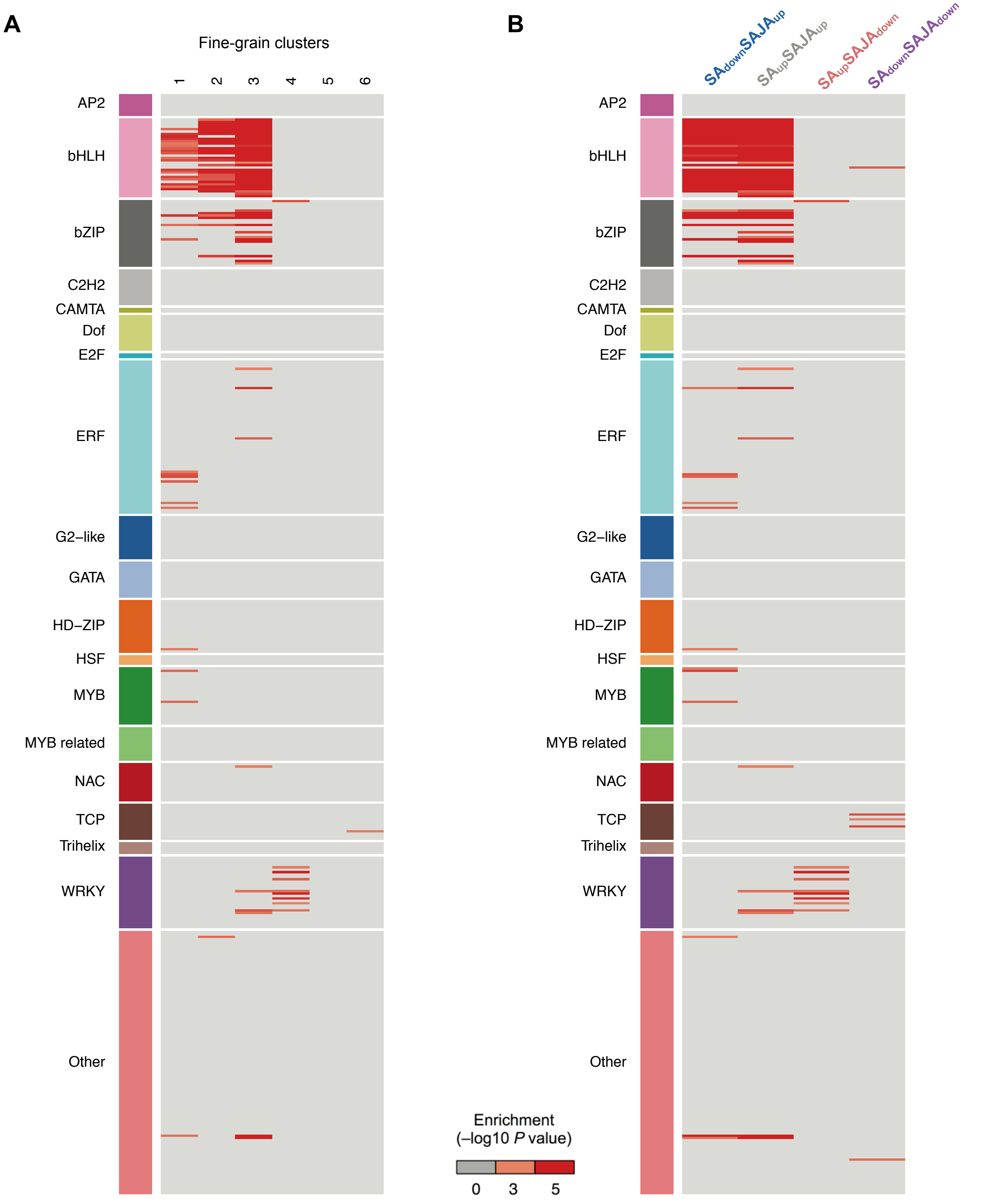


**Supplemental Figure S3. Enriched *cis*-regulatory motifs in clusters of SA-responsive genes that are sensitive to MeJA crosstalk.**
(Supports Figure 7D)

Known TF DNA-binding motifs enriched in the promoters of genes clustered based on their gene expression profiles responding to SA alone and to the combined SA- and MeJA treatment as shown in Figure 6D. **(A)** Motif enrichment *P* values in the 6 fine-grain clusters. **(B)** Motif enrichment *P* values in the course-grain clusters. Columns indicate clusters. Rows indicate motifs and are colored by corresponding TF family. Red boxes indicate a motif that is significantly overrepresented (hypergeometric test).

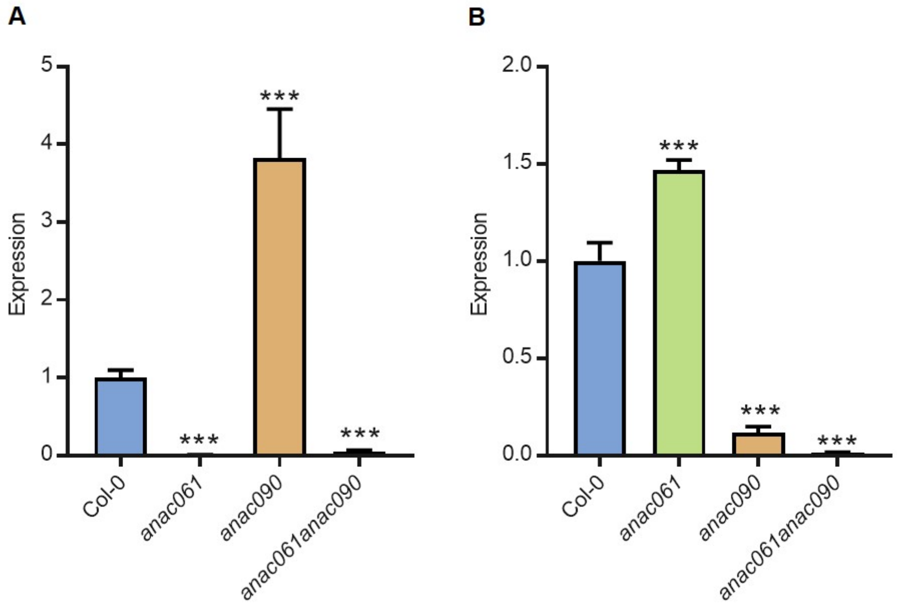


**Supplemental Figure S4. Expression of *ANAC061* and *ANAC090* in Col-0 (WT), *anac061*, *anac090* and *anac061 anac090.***
(Supports Figure 8)

Transcript abundance of *ANAC061* **(A)** and *ANAC090* **(B)** in single mutant insertion alleles and corresponding double mutant determined using qRT-PCR, 1 h after leaves were syringe-inoculated with 1 µM of flg22. Transcript abundance is expressed relative to the wild type (Col-0) and is the average and SE of three biological replicates. ****P* value < 0.0001.

| AGI | Gene name | Mutant | Cluster | *Pst* DC3000 phenotype |
|---|---|---|---|---|
| AT1G66560 | *WRKY64* | *wrky64* | 2 | NS |
| AT1G80590 | *WRKY66* | *wrky66* | 2 | NS |
| AT1G66550 | *WRKY67* | *wrky67* | 2 | NS |
| AT5G09330 | *ANAC082* | *anac082* | 5 | NS |
| AT1G57560 | *MYB50* | *myb50* | 5 | NS |
| AT1G17460 | *TRFL3* | *trfl3* | 6 | NS |
| AT3G04070 | *ANAC047* | *anac047* | 7 | NS |
| AT3G10500 | *ANAC053* | *anac053* | 7 | NS |
| AT1G09540 | *MYB61* | *myb61* | 10 | NS |
| AT4G36710 | *HAM4* | *ham4* | 13 | NS |
| AT5G04410 | *ANAC078* | *anac078* | 13 | NS |
| AT2G40950 | *bZIP17* | *bzip17* | 15 | NS |
| AT1G34190 | *ANAC017* | *anac017* | 15 | NS |
| AT5G63790 | *ANAC102* | *anac102* | 19 | NS |
| AT1G02230 | *ANAC004* | *anac004* | 20 | NS |
| AT3G44350 | *ANAC061* | *anac061* | 22 | NS |
| AT5G22380 | *ANAC090* | *anac090* | 22 | *RDS |
| AT5G64530 | *ANAC104* | *anac104* | 25 | NS |
| AT1G57560<br>AT1G09540 | *MYB50*<br>*MYB61* | *myb50 myb61* | 5,10 | NS |
| AT3G10500<br>AT5G04410 | *ANAC053*<br>*ANAC078* | *anac053 anac078* | 7,13 | NS |
| AT5G64530<br>AT5G22380 | *ANAC061*<br>*ANAC090* | *anac061 anac090* | 22,22 | **RDS |

**Supplementary Table 1. A list of all mutant alleles for predicted SA-pathway regulators and their *Pst* DC3000 disease resistance phenotypes.**

*P. syringae* DC3000 resistance phenotypes of TF mutant lines based on bacterial growth in leaves 72 h after pressure infiltration. Mutants were tested in different batches together with Col-0 (WT). One-way ANOVA with Dunnett's post-hoc test was used to compare the $\log_{10}$-transformed cfu/cm$^2$ of leaf tissue of the different genotypes to WT. *$P < 0.05$; **$P < 0.001$; NS, no significant difference; RDS, reduced disease symptoms.
